# Spatial rearrangement of the *Streptomyces venezuelae* linear chromosome during sporogenic development

**DOI:** 10.1101/2020.12.09.403915

**Authors:** MJ. Szafran, T. Małecki, A. Strzałka, K. Pawlikiewicz, J. Duława, A. Zarek, A. Kois-Ostrowska, K. Findlay, TBK. Le, D. Jakimowicz

## Abstract

Depending on the species, bacteria organize their chromosomes with either spatially separated or closely juxtaposed replichores. However, in contrast to eukaryotes, significant changes in bacterial chromosome conformation during the cell cycle have not been demonstrated to date. *Streptomyces* are unique among bacteria due to their linear chromosomes and complex life cycle. These bacteria develop multigenomic hyphae that differentiate into chains of unigenomic exospores. Only during sporulation-associated cell division, chromosomes are segregated and compacted. In this study, we show that at entry to sporulation, arms of *S. venezuelae* chromosomes are spatially separated, but they are closely aligned within the core region during sporogenic cell division. Arm juxtaposition is imposed by the segregation protein ParB and condensin SMC. Moreover, we disclose that the chromosomal terminal regions are organized into domains by the *Streptomyces*-specific protein - HupS. Thus, we demonstrate chromosomal rearrangement from open to close conformation during *Streptomyces* life cycle.

## INTRODUCTION

Chromosomal organization differs notably among the domains of life, and even among bacteria, chromosomal arrangement is not uniform. While linear eukaryotic chromosomes undergo dramatic condensation before their segregation, the compaction of usually circular bacterial chromosomes remains relatively constant throughout the cell cycle^1–3^. To date, chromosome conformation capture methods, applied in only a few model species, have revealed that bacterial chromosomes adopt one of the two distinct conformation patterns. Either both replichores of circular chromosomes align closely (as in *Bacillus subtilis*^4,5^, *Caulobacter crescentus*^6^, *Corynebacterium glutamicum*^7^, *Mycoplasma pneumoniae*^8^, and chromosome II of *Vibrio cholerae*^9^) or tend to adopt an “open” conformation, which lacks close contacts between both chromosomal arms (e.g., in *Escherichia coli*^10^ and *V. cholerae* chromosome I^9^). The cytological studies showed that chromosome organisation may alter depending on the growth phase or developmental stage^1,11,12,14^. While the most prominent change in bacterial chromosome organization was observed during extensive DNA compaction that occurs during sporulation of *Bacillus spp*^13^ and *Streptomyces spp*s, the details of chromosome arrangement during this process have not been elucidated.

The folding of bacterial genomes requires a strictly controlled interplay between several cellular factors that encompass: macromolecular crowding, transcription and the activities of dedicated DNA-organizing proteins - topoisomerases, condensins and nucleoid-associated proteins (NAPs)^10,15,16^. By nonspecific DNA binding, NAPs induce its bending, bridging or wrapping to govern the local organization and global compaction of bacterial chromosomes^1,10,16^. Since the subcellular repertoire of NAPs is dependent on growth conditions, changes in their supply account for the fluctuation of chromosome compaction in response to environmental clues. While, the repertoire of NAPs also varies between bacterial species, some of them, such as HU homologues, are widespread among bacteria^14,17,18^.

In contrast to NAPs, condensins are the most conserved DNA-organizing proteins identified in all kingdoms of life. In bacteria, the homologues of condensin are complexes of SMCs or its *E. coli* functional analogue MukB, associated with their accessory proteins (ScpAB or MukEF, respectively) ^19,20,21^. According to a recently proposed model, the primary activity of SMC is the extrusion of large DNA loops^22,23^. SMC complexes were shown to move along bacterial chromosomes from the centrally positioned origin of replication (*oriC*), leading to gradual chromosome compaction and parallel alignment of both replichores^5,6,24,25^. Importantly, although the alignment of two chromosomal arms was not detected in *E. coli*, the MukBEF complex is still responsible for the long-range DNA interactions within chromosomal domains^10,26^.

In *B. subtilis, C. glutamicum*, and *C. crescentus*, the loading of SMC in the vicinity of the *oriC* was shown to be dependent on the segregation protein ParB^4,27,28^. Binding to a number of *parS* sequences, ParB forms a large nucleoprotein complex (segrosome), that interact with the partner protein ParA, ATPase non-specifically bound to nucleoid ^29,30^. ParB interaction with ParA triggers its ATP hydrolysis and nucleoid release, generating a concentration gradient that drives segrosome separation^31–33^. Segrosome formation was shown to be a prerequisite for chromosomal arm alignment in the model Gram-positive bacteria with circular chromosomes^5,7,28^. Importantly, there have been no reports on the 3D organization of bacterial linear chromosomes to date.

The example bacteria with linear chromosomes are *Streptomyces* - Gram-positive mycelial soil actinobacteria, which are highly valued producers of numerous antibiotics and other clinically important metabolites. Notably, the production of these secondary metabolites is closely correlated with *Streptomyces* unique and complex life cycle, which encompasses vegetative growth and sporulation, as well as exploratory growth in some species^14,34,35^. During the vegetative stage, *Streptomyces* grow as branched hyphae composed of elongated cell compartments, each containing multiple copies of nonseparated and largely uncondensed chromosomes^14,36^. Under stress conditions sporogenic hyphae develop, and their rapid elongation is accompanied by extensive replication of chromosomes^37^. When hyphae stop extending, the multiple chromosomes undergo massive compaction and segregation to unigenomic spores^14^. These processes are accompanied by multiple synchronized cell divisions initiated by the FtsZ protein, which assembles into a ladder of regularly spaced Z-rings^38,39^. Chromosome segregation is driven by ParA and ParB proteins, whereas nucleoid compaction is induced by concerted action of SMC and sporulation-specific NAPs that include HupS - one of the two *Streptomyces* HU homologues, structurally unique to actinobacteria due to the presence of histone-like domain. The elimination of any of these proteins results in DNA decompaction in spores and affects resistance to environmental factors^40–44^. Thus, while in unicellular bacteria chromosome segregation occurs during ongoing replication, during *Streptomyces* sporulation, intensive chromosome replication is temporarily separated from their compaction and segregation, with the two latter processes occurring only during sporogenic cell division.

While the *Streptomyces* chromosomes are among the largest in bacterial kingdom, the knowledge concerning their organization is scarce. Limited cytological evidence suggests that during vegetative growth, both ends of the linear chromosome colocalize, and the *oriC* region of the apical chromosome is positioned at the tip-proximal edge of the nucleoid^45,46^. The visualization of nucleoid and ParB-marked *oriC* regions in sporogenic hyphae showed that *oriC*s are positioned centrally within pre-spore compartments, but did not elucidate their architecture during sporulation^39,46^.

In this study, using the chromosome conformation capture method (Hi-C), we investigated *S. venezuelae* chromosome organization during differentiation. The application of the Hi-C technique demonstrated the dramatic rearrangement of chromosome structure associated with sporogenic development. Moreover, by combining the Hi-C method with chromatin immunoprecipitation and fluorescence microscopy techniques, we showed the contribution of ParB, SMC and HupS (the sporulation-dedicated HU homologue) to global nucleoid compaction.

## MATERIALS and METHODS

### Bacterial strains and plasmids

The *S. venezuelae* and *E. coli* strains utilized in the study are listed in Table S1 and S2 respectively, and the plasmids are listed in Table S3. DNA manipulations were performed by standard protocols^47^. Culture conditions, antibiotic concentrations, transformation and conjugation methods followed the commonly employed procedures for *E. coli*^47^ and *Streptomyces*^48^. A detailed description of strain construction is presented in the Supplementary Information.

### *S. venezuelae* cultures and growth rate analysis

To obtain the standard growth curves comparing the growth rate of *S. venezuelae* strains, a Bioscreen C instrument (Growth Curves US) was used. Cultures (in triplicate) in liquid MYM medium (300 µl) were set up by inoculation with 6 × 10^3^ colony-forming units (CFUs) of *S. venezuelae* spores. The cultures grew for 48 h at 30°C under the “medium” speed and shaking amplitude settings, and their growth was monitored by optical density measurement (OD_600_) every 20 min.

The Hi-C cultures were set up as follows: 5 ml of MYM medium was inoculated with 10^8^ CFU, and the cultures were incubated for 6 to 26 h at 30°C with shaking (180 rpm). For the “5 ml culture” growth curve depiction, the optical density (OD_600_) of the culture diluted (1:10) with MYM medium was measured in 1-h.

### Preparation of *Hi-C* libraries and data analysis

For preparation of *S. venezuelae* Hi-C contact maps, 5 ml cultures were established as described above. The cultures were incubated for 13 to 25 h at 30°C with shaking (180 rpm). After this step, the cultures were cross-linked with 1% formaldehyde for 7 min and 30 s and blocked with 0.2 M glycine for 15 min. One millilitre of the cross-linked culture was centrifuged (1 min, 5000 rpm at room temperature). The mycelium pellet was washed twice with 1.2 ml of TE buffer and resuspended in 0.6 ml of TE buffer. Two vials, each containing 25 µl of the resuspended mycelium, were independently processed according to the modified protocol of Le *et al*.^49^ as below. Briefly, the cells were lysed using 0.5 µl of ReadyLyse solution (Invitrogen) at 37°C for 30 min followed by the addition of 1.25 µl of 5% SDS solution and incubation at room temperature for 15 min. After cell lysis, the reaction was supplemented with 5 µl of 10% Triton X-100 (Sigma Aldrich), 5 µl of NEB3 buffer (New England Biolabs) and 10.75 µl of DNase-free water (Invitrogen) and subsequently incubated for 15 min at room temperature. The chromosomal DNA was digested with 2.25 µl of *BglII* restriction enzyme (New England Biolabs; 50000 U/ml) for 2 h and 30 min at 37°C followed by the addition of 0.25 µl of *BglII* and incubation for another 30 min at 37°C. The subsequent steps of Hi-C library preparation directly followed the protocol described by Le *et al*.^49^with the number of amplification cycles being increased to 18. The amplified libraries were purified from a 1% agarose gel, and their concentrations were measured using a Qubit dsDNA HS Assay Kit (Thermo Fisher Scientific) in OptiPlate-96 White (Perkin Elmer) with 490-nm and 520-nm excitation and emission wavelengths, respectively. The agarose-purified Hi-C libraries were diluted to a 10 nM concentration, pooled together, and sequenced using Illumina NextSeq500 (2×75 bp; paired reads), as offered by the Fasteris SA sequencing facility.

Complete data processing was performed using the Galaxy HiCExplorer web server^50^. The paired reads were first trimmed using *Trimmomatic* (version 0.36.5) and subsequently filtered using *PRINSEQ* (version 0.20.4) to remove reads with more than 90% GC content. The pre-processed reads were mapped independently to the *S. venezuelae* chromosome using *Bowtie2* (version 2.3.4.3; with the -*sensitive-local* presets). The Hi-C matrix was generated using *hicBuildMatrix* (version 2.1.4.0) with a bin size of 10000 bp. In the next step, three neighbouring bins were merged using *hicMergeMatrixBins* (version 3.3.1.0) and corrected using the *hicCorrectMatrix* tool (version 3.3.1.0) based on Imakaev’s iterative correction algorithm^51^ (with the presets as follows: number of iterations: *500*; skip diagonal counts: *false*; remove bins of low coverage: *-1*.*5*; remove bins of high coverage: *5*.*0*). The corrected Hi-C matrixes were normalized and compared using *hiCCompareMatrixes* (version 3.4.3.0, presets as below: *log2ratio)*, yielding differential Hi-C contact maps. The differential maps were plotted using *hicPlotMatrix* (version 3.4.3.0; with masked bin removal and *seismic* pallets of colours). To prepare the Hi-C contact map, the corrected matrixes were normalized in the 0 to 1 range using *hicNormalize* (version 3.4.3.0; settings: set values below the threshold to zero: *value 0*) and subsequently plotted using *hicPlotMatrix* (version 3.4.3.0; with the removal of the masked bins), yielding 30-kb-resolution Hi-C heatmaps.

### Chromatin immunoprecipitation combined with next-generation sequencing (Chip-seq)

For chromatin immunoprecipitation, *S. venezuelae* cultures (in triplicate) were grown in 50 ml liquid MYM medium supplemented with TES at 30°C with shaking (ChIP-seq cultures). Cultures were inoculated with 7×10^8^ CFU. After 14 h of growth, the cultures were cross-linked with 1% formaldehyde for 30 min and blocked with 125 mM glycine. Next, the cultures were washed twice with PBS buffer, and the pellet obtained from half of the culture volume was resuspended in 750 μl of lysis buffer (10 mM Tris-HCl, pH 8.0, 50 mM NaCl, 14 mg/ml lysozyme, protease inhibitor (Pierce)) and incubated at 37°C for 1 h. Next, 200 μl of zircon beads (0.1 mm, BioSpec products) were added to the samples, which were further disrupted using a FastPrep-24 Classic Instrument (MP Biomedicals; 2 x 45 s cycles at 6 m/s speed with 5-min breaks, during which the lysates were incubated on ice). Next, 750 μl IP buffer (50 mM Tris-HCl pH 8.0, 250 mM NaCl, 0.8% Triton X-100, protease inhibitor (Pierce)) was added, and the samples were sonicated to shear DNA into fragments ranging from 300 to 500 bp (verified by agarose gel electrophoresis). The samples were centrifuged, 25 μl of the supernatant was stored to be used as control samples (“input”), and the remaining supernatant was used for chromatin immunoprecipitation and mixed with magnetic beads (60 μl) coated with anti-FLAG M2 antibody (Merck). Immunoprecipitation was performed overnight at 4°C. Next, the magnetic beads were washed twice with IP buffer, once with IP2 buffer (50 mM Tris-HCl pH 8.0, 500 mM NaCl, 0.8% Triton X-100, protease inhibitors (Pierce)), and once with TE buffer (Tris-HCl, pH 7.6, 10 mM EDTA 10 mM). DNA was released by overnight incubation of beads resuspended in IP elution buffer (50 mM Tris-HCl pH 7.6, 10 mM EDTA, 1% SDS) at 65°C. The IP elution buffer was also added to the “input” samples, which were treated further as immunoprecipitated samples. Next, the samples were centrifuged, and proteinase K (Roche) was added to the supernatants to a final concentration of 100 µg/ml followed by incubation for 90 min at 55°C. DNA was extracted with phenol and chloroform and subsequently precipitated overnight with ethanol. The precipitated DNA was dissolved in nuclease-free water (10 µl). The concentration of the DNA was quantified using a Qubit dsDNA HS Assay Kit (Thermo Fisher Scientific).

DNA sequencing was performed by Fasteris SA (Switzerland) using the Illumina ChIP-Seq TruSeq protocol, which included quality control, library preparation and sequencing from both ends (2×75 bp) of amplified fragments. The bioinformatic analysis was performed using the R packages *edgeR, normr, rGADEM*, and *csaw*, as well as the *MACS2* and *MEME* programs. HupS-FLAG binding was analysed using an earlier published protocol^52^. The mapping of ChIP-seq data was performed using the *Bowtie2* tool (version 2.3.5.1^53,54^). The successfully mapped reads were subsequently sorted using *samtools* (version 1.10)^55^. The total number of mapped reads was above 10^6^ on average. Regions differentially bound by HupS-FLAG were identified using the R packages *csaw* and *edgeR*^56–59^, which also normalizes the data, as described earlier by Lun and Smyth^52^. The mapped reads were counted within a 69-bp-long sliding window with a 23-bp slide. Peaks were filtered using the local method from the *edgeR* package, which compares the number of reads in the region to the background computed in the surrounding window of 2000 bp. Only regions with log2 fold change above 2.0 were utilized for further analysis. Regions that were less than 100 bp apart were merged, and the combined p-value for each was calculated. Only regions with false discovery rate (FDR) values below the 0.05 threshold were considered to be differentially bound. The identified regions were further confirmed by analysis with the *MASC2* program using the *broad* option^60^. The R packages *rGADEM* and *MEME* Suite were used to search for probable HupS binding sites^61^. SMC-FLAG bound regions were analysed using the R package *normr* using Input ChIP-seq data as a control. Reads were counted in 250-bp-long regions, and a region was considered to be bound by SMC-FLAG if FDR value compared to Input was below the 0.05 threshold, and if it was not found to be significant in a control strain not producing SMC-FLAG protein^62^.

### Quantification of the *oriC/ter* ratio

To estimate the *oriC/ter* ratio, chromosomal DNA was extracted from *S. venezuelae* 5 ml cultures (as for Hi-C) growing for 13-26 h or from 50 ml culture (as for ChIP-seq) growing for 8 to 16 h. One millilitre of each culture was centrifuged (1 min at 5000 rpm and at room temperature). The supernatant was discarded, and the mycelium was used for the subsequent isolation of chromosomal DNA with application a Genomic Mini AX *Streptomyces* kit (A&A Biotechnology) according to the manufacturer’s protocol. The purified DNA was dissolved in 50 µl of DNase-free water (Invitrogen). The DNA concentration was measured at 260 nm and subsequently diluted to a final concentration of 1 ng/ml.

qPCR was performed using Power Up SYBR Green Master Mix (Applied Biosystems) with 2 ng of chromosomal DNA serving as a template for the reaction and oligonucleotides complemented to the *oriC* region (*oriC*) (gyr2_Fd, gyr2_Rv) or to the termination region (*ter*) (arg3_Fd, arg3_Rv)(Table S4). The changes in the *oriC/ter* ratio were calculated using the comparative ΔΔCt method, with the *ter* region being set as the endogenous control. The *ori*/*ter* ratio was estimated as 1 in the culture (5 ml) growing for 26 h, corresponding to the appearance of spore chains.

### Light Microscopy

The analyses of spore sizes were performed using brightfield microscopy. Spores from 72-h plate cultures (MYM medium without antibiotics) were transferred to microscopy slides. The samples were observed using a Zeiss Microscope (Axio Imager M1), and images were analysed using Fiji software. The lengths of 300 spores were measured for each strain.

For the snapshot analysis of *S. venezuelae* development in liquid, 5 µl from the 5 ml culture (as for Hi-C) growing for 18 to 26 h or from the 50-ml culture (as for ChIP-Seq) growing for 8 to16 h was spread on a coverslip and fixed by washing twice with absolute methanol. The nucleoids were subsequently stained for 5 min at room temperature with 7-amino-actinomycin D (1 mg/ml 7-AAD in DMSO, Thermo Fisher Scientific) diluted 1:400 in PBS buffer. In the next step, the mycelium was washed twice with PBS buffer, and the coverslip was mounted using 50% glycerol in PBS. Microscopic observation of FtsZ-YPET and DNA stained with 7-AAD was performed using a Zeiss Microscope (Axio Imager M1). The images were analysed with dedicated software (Axio Vision Rel software). To calculate the nucleoid separation, the distance between the two neighbouring DNA-free areas was measured for 250 nucleoids. The boxplots showing the length of nucleoid-covered areas were constructed using the Plotly tool (plotly.com).

Time-lapse fluorescence microscopy was performed using a previously described protocol^38^ with modification of the applied pressure (3 psi), increased dilution of the loaded spores, and application of repeated spore loading. The observation was performed using a DeltaVision Elite Ultra High-Resolution microscope equipped with a camera and the CellAsic Onix microfluidic system using B04A plates (ONIX). The images were taken every 10 min for approximately 20 h using DV Elite CoolSnap and 100X, 1.46 NA, DIC lens for specimens exposed for 50 ms at 5% transmission in the DIC channel and for 80 ms at 50% transmission in the YFP and mCherry channels. The time-lapse movies were analysed using ImageJ Fiji software. Nucleoid area was measured using HupA-mCherry as the marker at the time of disassembly of Z-rings (visualized using FtsZ-Ypet fusion), which was approximately 120 min after their appearance and hyphal growth cessation. Analysis of the HupA-mCherry-labelled chromosome images was based on determining the area and Feret diameter (the maximal distance between two points) of nucleoid fluorescence using the Versatile Wand Tool with 8-connected mode and manually adjusted value tolerance. The distances between Z-rings were measured 60 min after hyphal growth cessation for approximately 300 Z-rings. The fluorescence intensity of the Z-rings was collected along and averaged across the hyphae with a line selection tool whose width was adjusted to the width of the measured hyphae. The fluorescence intensity of each Z-ring in 25-30 hyphae of each strain was subsequently determined by analysing the data with RStudio using a custom shiny application *findpeaks* based on the *Peaks* R package. Tracking of Z-rings in the time-lapse images was performed using the nearest neighbour algorithm. Only Z-rings observed for more than 90 min were used in the analysis. The data from each time point were combined to show how the standard deviation of fluorescence intensity changed during hyphal development. Representative hyphae of each strain were selected, the FtsZ-YPet fluorescence intensity and Z-ring position were determined using the script described above, and its variability was visualized in the form of a kymograph with a custom script based on the *ggplot2* package.

### Scanning Electron Microscopy

*Streptomyces* samples were mounted on an aluminium stub using Tissue Tek^R^ OCT (optimal cutting temperature) compound (BDH Laboratory Supplies, Poole, England). The stub was then immediately plunged into liquid nitrogen slush at approximately −210°C to cryo-preserve the material. The sample was transferred onto the cryo-stage of an ALTO 2500 cryo-transfer system (Gatan, Oxford, UK) attached to an FEI Nova NanoSEM 450 (FEI, Eindhoven, The Netherlands). Sublimation of surface frost was performed at −95°C for approximately three minutes before sputter coating the sample with platinum for 150 seconds at 10 mA, at colder than −110°C. After sputter-coating, the sample was moved onto the cryo-stage in the main chamber of the microscope, held at −125°C. The sample was imaged at 3 kV and digital TIFF files were stored.

## RESULTS

### Hi-C reveals an alignment chromosomal arms extending to terminal domains during *S. venezuelae* sporogenic cell division

During *Streptomyces* sporulation, their multigenomic hyphae undergo considerable morphological changes turning into a chains of spores through multiple cell divisions. This event is linked to inhibition of DNA replication followed by synchronized compaction and segregation of a dozen chromosomes^38,39,64–66^. To explore whether the changes in chromosome compaction during sporulation are reflected in its global organization, we applied the chromosome conformation capture technique (Hi-C), which enabled us to establish a map of DNA contact frequency^6,67,68^.

First, we determined the time points of critical sporulation events for *S. venezuelae*, namely, the reduction of hyphal growth and cessation of DNA replication, chromosome compaction and separation, formation of the FtsZ ladder, and appearance of spore chains (**Fig. 1A** and **S1**). After 13-14 h of growth (under 5 ml culture conditions), we observed vegetative growth inhibition. The slowed hyphal growth correlated with the reduction in the *oriC*/*ter* ratio corresponding to decreased DNA replication (**Fig. 1A**). These events indicated culture entry into a developmental stage that we named the pre-sporulation phase. During this phase, which lasted 5 h, the DNA replication rate transiently increased (at the 19^th^-21^st^ hour). This increased replication rate was observed during a slight increase in the optical density of the culture, which could be explained by the rapid growth of sporogenic hyphae (**Fig. 1A**). The subsequent drop in the replication rate was accompanied by an increase in the FtsZ-YPet level (at the 21^st^-23^rd^ hour), assembly of FtsZ-YPet into ladders of Z-rings (detected at the 22^nd^ hour), and initiation of chromosome compaction (**Figs. 1A** and **S1**). At this time point, the spore chains entered the maturation phase, during which chromosome condensation and separation increased, while the FtsZ-YPet level rapidly decreased and the hyphae fragmented. This stage ended when mature spores could be detected (at the 25^th^-26^th^ hour) (**Fig. 1A** and **Fig. S1**). Thus, we established that 22^nd^ hour of *S. venezuelae* culture is the time point when multigenomic hyphae undergo synchronized cell division, replication of chromosomes ceases, while their condensation increases, which minimizes interchromosomal contacts. Based on these findings, we set out to establish the chromosome conformation at this time point.

**Figure 1.**
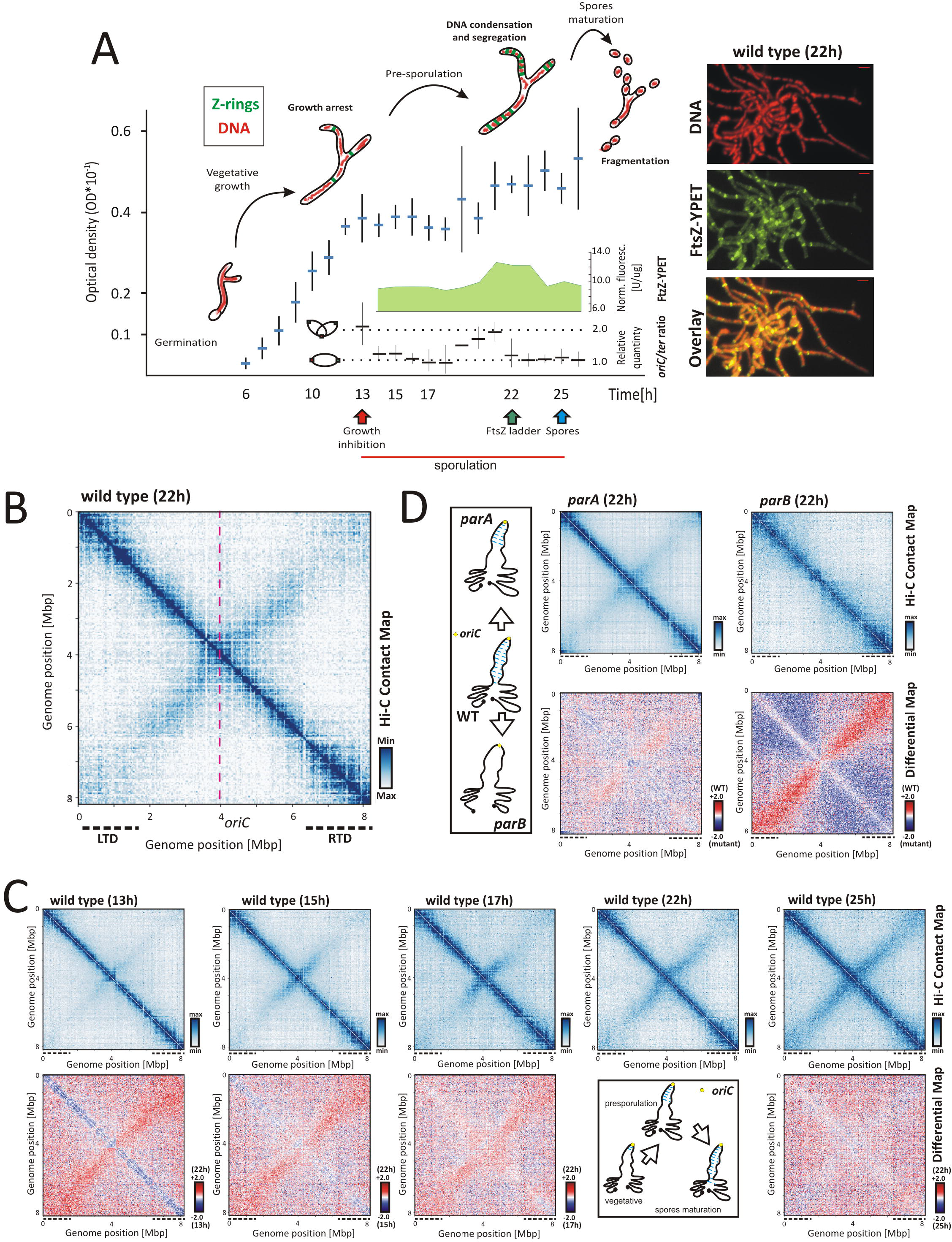
Spatial organization of the *S. venezuelae* linear chromosome during sporulation. The *S. venezuelae* (wild type *ftsZ-ypet* derivative, MD100) growth curve (in 5 ml culture). The critical time points of sporogenic development are marked with arrows: growth arrest (red), the appearance of FtsZ ladders (green) and the formation of spore chains (blue). The normalized FtsZ-YPet fluorescence [U/µg] and the relative *oriC*/*ter* ratio are shown as insets in the plot, with X axes corresponding to the main plot X axis. The *oriC/ter* ratio at the 25^th^ hour of growth was set as 1.0. The right panel shows the visualization of condensing nucleoids (DNA with staining 7-AAD) and FtsZ-YPet at 22^nd^ hour of growth; scale bar - 2 µm. The normalized Hi-C contact map obtained for the wild type (*ftsZ-ypet* derivative, MD100) after 22 h of growth (5 ml culture). The dotted pink lines mark the position of the *oriC* region. The dotted black lines mark the positions of the left (LTD) and right (RTD) terminal domains. (**C**) Top panel: the normalized Hi-C contact maps obtained for the wild type (*ftsZ-ypet* derivative, MD100) strain growing for 13, 15, 17, 22 and 25 h (in 5 ml culture). Bottom panel: the differential Hi-C maps in the logarithmic scale (log2) comparing the contact enrichment at 22 h of growth (red) versus other time points (blue). The X- and Y-axes indicate chromosomal coordinates binned in 30 kb. The inset shows the model of chromosome rearrangement in the course of sporulation. (**D**) Top panel: the normalized Hi-C contact maps generated for *parA* and *parB* mutants (*ftsZ-ypet* derivatives MD011 and MD021, respectively) growing for 22 h (5 ml culture). Bottom panel: the differential Hi-C maps in the logarithmic scale (log2) comparing the contact enrichment in the wild type strain (red) versus the mutant strain (blue). The inset shows the model of chromosome organization in the *parA* and *parB* mutants.

Hi-C mapping of the chromosomal contacts in *S. venezuelae* sporogenic hyphae undergoing cell division indicated the presence of two diagonal axes crossing perpendicularly in the proximity of the centrally located *oriC* region (**Fig. 1B**). The primary diagonal corresponds to the contact frequency between neighbouring chromosomal loci. We distinguished (based on PCA1/PCA2 analysis) three distinct chromosome regions - the core region that encompasses 4.4 Mbp (position: 1.9 Mbp to 6.3 Mbp) around the centrally positioned *oriC* and two well-organized domains flanking the chromosome core, which we named the left and right terminal domains (LTD and RTD, respectively), with each measuring approximately 2 Mbp in size (**Fig. 1B** and **S2A**). Interestingly, the boundaries of LTD and, especially, RTD overlap the chromosomal regions with significantly lower transcriptional activity in comparison to the chromosome core region (**Fig. S2B**), suggesting a correlation between chromosome organization and transcription. The secondary diagonal, that results from the interarm interactions, was not complete during cell divisions and was limited in each direction to approximately 2.0-2.2 Mbp from the *oriC* region (**Fig. 1B**). Additionally, the Hi-C contact map showed signals at the ends of the secondary diagonal, suggesting the existence of the spatial juxtaposition of terminal ends of the linear *S. venezuelae* chromosome (**Fig. 1B**).

In summary, our results demonstrated that at the time of *S. venezuelae* synchronized cell division, the linear chromosomes are arranged with both arms closely aligned within the core region, while the terminal regions form distinct domains.

### Arm juxtaposition progresses during sporulation and requires segregation protein activity

Having established chromosomal contact maps during *S. venezuelae* sporogenic cell division, we set out to characterize the changes in chromosomal arrangement during sporogenic development. To this end, we obtained normalized Hi-C contact maps for the wild type strain at the entry into the pre-sporulation phase when vegetative growth and DNA replication decrease (at the 13^th^ hour) and during the phase when the chromosomes are not fully compacted (at the 15^th^ and 17^th^ hours), as well as during at the final stage of spore maturation, which is when chromosome compaction reaches a maximum (**Fig. 1A** and **S1**).

At the entry into the pre-sporulation phase (at the 13^th^ hour), only the primary diagonal was clearly visible, whereas the secondary diagonal was barely detectable (**Fig. 1C)**. Additionally, the juxtaposition of chromosomal termini was also not, or only slightly, noticeable at this stage of growth. During the pre-sporulation phase, the signals of the secondary diagonal were enhanced and extended at distances of up to 1 Mbp from the *oriC* region at the 15^th^ and 17^th^ hours of growth (**Fig. 1C**). Finally, in the mature spores (at the 25^th^ hour), we observed almost complete arm alignment, similar to that detected at 22^nd^ hour (**Fig. 1C**). This result suggests that although nucleoid compaction was still underway until the end of the spores maturation phase (**Fig. S2B**), the global chromosome organization did not change critically after that established during cell division (**Fig. 1C**).

In model bacteria whose chromosomal arms are juxtaposed, their close positioning is imposed by ParB-dependent SMC loading within the *oriC* region^4,7,28^. During *Streptomyces* sporogenic cell division, at the time of observed chromosomal arm juxtaposition, ParB was shown to assemble into regularly spaced complexes that position *oriC* regions along the hyphae^69^. At the same time, ParA accumulates in sporogenic hyphae and facilitates the efficient formation and segregation of ParB complexes^39,70^. The elimination of *S. venezuelae* segregation proteins disturbed chromosome segregation^39^ but did not affect the growth rate (**Fig. S5A**). To verify whether segregation proteins play a role in the arrangement of *S. venezuelae* chromosomes, we obtained Hi-C maps of chromosomal contacts in sporogenic hyphae of *parA* or *parB* mutants.

The Hi-C map generated for the *parB* mutant showed complete disappearance of the secondary diagonal axis, reinforcing the role of the ParB complex in inducing interarm contacts (**Fig. 1D**). The same lack of secondary diagonal was observed for the *parB* mutant at each time of *S. venezuelae* sporulation (**Fig. S3B)**. The influence of the *parA* deletion on arm alignment was also detectable, although it varied among the samples analysed (**Fig. 1C** and **S3C**). The strength of the signals at the diagonal axis was either significantly or only slightly lowered in the *parA* mutant, as indicated by the differential Hi-C map in comparison to the wild type strain (**Fig. 1C** and **S3C**). While the effect of *parA* or *parB* deletions was predominantly manifested by the partial or complete disappearance of the interarm contacts, respectively, their influence on the local chromosome structure, including LTD and RTD folding, was marginal (**Fig. 1C** and **S3**).

In summary, these results indicated that chromosomal arm alignment progressed during pre-sporogenic and sporogenic hyphal development (**Fig. 1C, inset**) and chromosomal rearrangement was dependent on ParB.

### SMC- and HupS-induced chromosome compaction permits regularly spaced cell division

In numerous bacteria, the role of ParB in chromosome organization is associated with SMC loading which, on the other hand, promotes global chromosome compaction. Prior *S. coelicolor* studies showed that elimination of either SMC- or *Streptomyces*-specific sporulation-associated NAP-HupS affected nucleoid compaction in spores^40,42,71^. We predicted that SMC would be responsible for ParB-induced chromosomal arm alignment, but since ParB elimination did not influence short-range chromosome interactions, we hypothesized that other DNA organizing factors, such as HupS, may also contribute to chromosome compaction in sporogenic hyphae. Therefore, in this study, we analysed sporulation and chromosome organization in *hupS* and *smc* mutant strains of *S. venezuelae*.

The disruption of either the *hupS* or *smc* gene in *S. venezuelae* did not significantly affect the growth rate or differentiation of these bacteria; however, the double *hupS smc* mutant growth rate was slower than that of single mutants (**Fig. S4A, inset** and **S4B**). Notably, the elimination of *hupS* or, to a lesser extent, *smc* increased spore size, and the double *hupS smc* deletion led to more pronounced disturbances in spore size than single deletion (**Fig. 2A and 2B**). The *hupS* or *smc* deletion phenotype, manifested by spore length, was restored by complementation of deletion strains with *in trans*-delivered *hupS*-*FLAG* or *smc-FLAG* genes, respectively (**Fig. 2B**).

**Figure 2.**
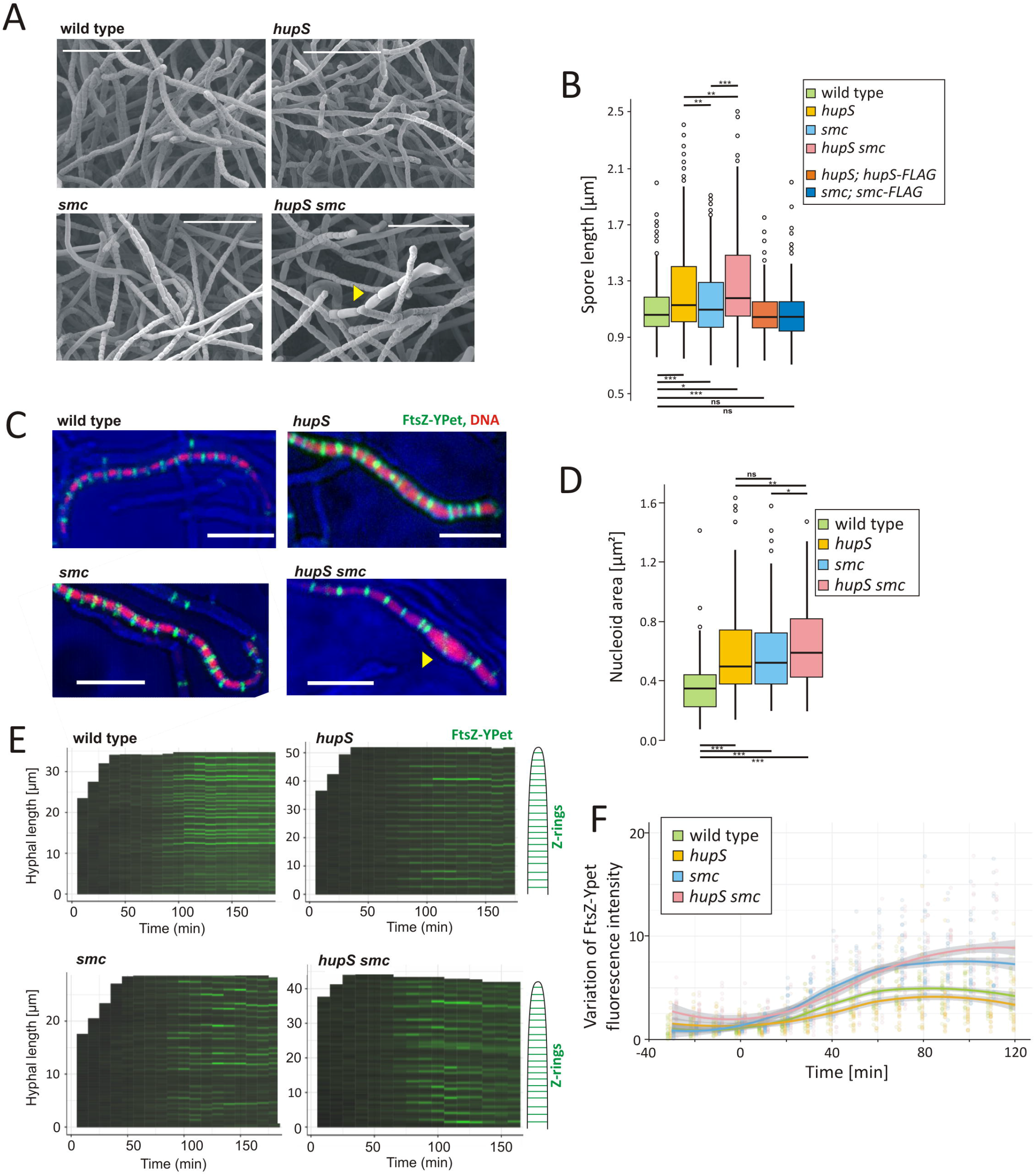
Phenotypic effects of *smc, hupS* and double *smc/hupS* deletions. **(A)** Scanning electron micrographs showing sporulating hyphae and spores of the wild type and *hupS, smc* and *hupS smc* double mutant (*ftsZ-ypet* derivatives, MD100, TM005, TM004 and TM006, respectively). Elongated spores are marked with a yellow arrowhead. Scale bar: 10 µm. (**B)** Box plot analysis of the spore length distribution in the *S. venezuelae* wild type (light green), *hupS* (yellow), *smc* (blue) and *hupS smc* (pink) (AK200, TM010, TM003, respectively), as well as in *hupS* and *smc* mutants complemented with *in trans* delivered *hupS-flag* (orange) and *smc-flag* genes (dark blue) (TM015 and TM019, respectively) (approximately 300 spores were analysed for each strain). The statistical significance between strains is marked with asterisks. **(C)** Representative examples of time-lapse images showing visualization of nucleoids (marked with mCherry-HupA fusion) and Z-rings (visualized by FtsZ-YPet fusion) in the wild type background (*ftsZ-ypet, hupA-mCherry* derivative, TM011) *hupS, smc* and *hupS smc* double mutant background (*ftsZ-ypet, hupA-mCherry* derivative, TM013, TM012, and TM014, respectively). An abnormal hyphal fragment is marked with a yellow arrowhead. Scale bar: 5 µm. **(D)** Box plot analysis of the nucleoid area (visualized by mCherry-HupA fusion) in the wild type (light green), *hupS* (yellow), *smc* (blue) and *hupS smc* (pink) double mutant (*ftsZ-ypet, hupA-mcherry* derivative, MD100, TM011, TM013, TM012, TM014, respectively) **(E)** Kymographs showing the intensity of Z-ring fluorescence along the representative hyphae in the wild type and *hupS, smc* and *hupS smc* double mutant (*ftsZ-ypet* derivatives, MD100, TM005, TM004, TM006, respectively) during maturation **(F)** The variation (standard deviation) of FtsZ-Ypet (Z-rings) fluorescence during spore maturation of strains calculated for 25-30 hyphae of the wild type (light green), *hupS* (yellow), *smc* (blue) and *hupS smc* (pink) double mutant (*ftsZ-ypet* derivatives, TM005, TM004, TM006, respectively). Fluorescence was measured starting from 30 min before growth arrest until spore formation. Points represent values for each hyphal, while lines show loess model fit with 95% confidence intervals.

Next, to visualize nucleoid compaction and segregation during cell division, we observed sporogenic hyphal development of *smc* and/or *hupS* mutant strains labelled using HupA-mCherry and FtsZ-YPet (**Fig. 2C** and **S4C, Supplementary Movies 1-4**). None of the mutant strains formed anucleate spores, but all of them showed significantly increased nucleoid areas, and this phenotype was the most pronounced in the double *hupS smc* mutant, where notably enlarged spores could be observed (**Fig. 2A, C** and **D**, yellow arrowhead). The increased nucleoid area was accompanied by increased distances between Z-rings (**Fig. S4D**). The fraction of elongated (longer than 1.3 μm) pre-spore compartments was the highest in the double *hupS smc* mutant and in the *smc* mutant **(Fig. S4D)**. Additionally, the intensity of Z-rings in single *smc* or the double mutant was more varied along the hyphae than in the wild type strain, indicating their lowered stability (**Fig. 2E** and **F**). The disturbed positioning and stability of Z-rings explains the presence of elongated spores in mutant strains.

In summary, both SMC and HupS contribute to chromosome compaction in spores. The decreased chromosome compaction in the mutant strains correlated with decreased Z-ring stability and aberrant positioning which, in all studied mutants, resulted in increased distance between septa and elongated spores.

### SMC and HupS collaborate during sporogenic chromosome rearrangement

Having confirmed that both SMC and HupS mediate global chromosome compaction during *S. venezuelae* sporulation, we investigated how both proteins contribute to chromosomal contacts during sporulation-associated chromosome rearrangement.

The Hi-C contact map obtained for the *smc* mutant at sporogenic cell division (at the 22^nd^ hour) showed the complete disappearance of the secondary diagonal (similar to the map obtained for the *parB* mutant), indicating abolished chromosomal arm interactions (**Fig. 1B** and **3A)**. On the other hand, the elimination of *smc* resulted in a slight increase in short-range DNA contacts along the primary diagonal, whereas in the *parB* mutant, the short-range DNA contacts were not affected (**Fig. 1B, 3A** and **S5**).

Next, to confirm that ParB promotes the interaction of SMC with DNA, we analysed FLAG-SMC binding in the wild type background and in the *parB* mutant. The specific developmental time points of ChIP-seq experiment cultures (50-ml culture conditions) were identified, corresponding to the earlier described time points used for the Hi-C experiments (**Fig. 1A** and **S6A**). At the 12^th^ hour of “ChIP-seq culture”, DNA replication decreased (corresponding to the 15^th^-17^th^ hour of “Hi-C culture”), while at the 14^th^ hour of “ChIP-seq culture”, DNA compaction and the formation of regularly spaced Z-rings were clearly detected (**Fig. S6B**) (corresponding to the 22^nd^ hour in “Hi-C culture” conditions), indicating sporogenic cell division and chromosome segregation. Notably, the level of FLAG-SMC remind constant during sporogenic development (**Fig. S9C**).

The analysis of DNA fragments bound by FLAG-SMC at the time of sporogenic cell division did not identify any specific binding sites for FLAG-SMC, suggesting a lack of sequence preference. However, the quantification of the detected SMC binding sites along the chromosome showed that their frequency increased up to twofold in the chromosomal core region (**Fig. 3B** and **S7A**). The increased frequency of SMC binding overlapped with the secondary diagonal identified in the Hi-C experiments (**Fig. 3A, 3B** and **S7B**). SMC binding was severely diminished in the *parB* mutant (**Fig. 3B**), demonstrating that the ParB complex is a prerequisite for SMC loading.

**Figure 3.**
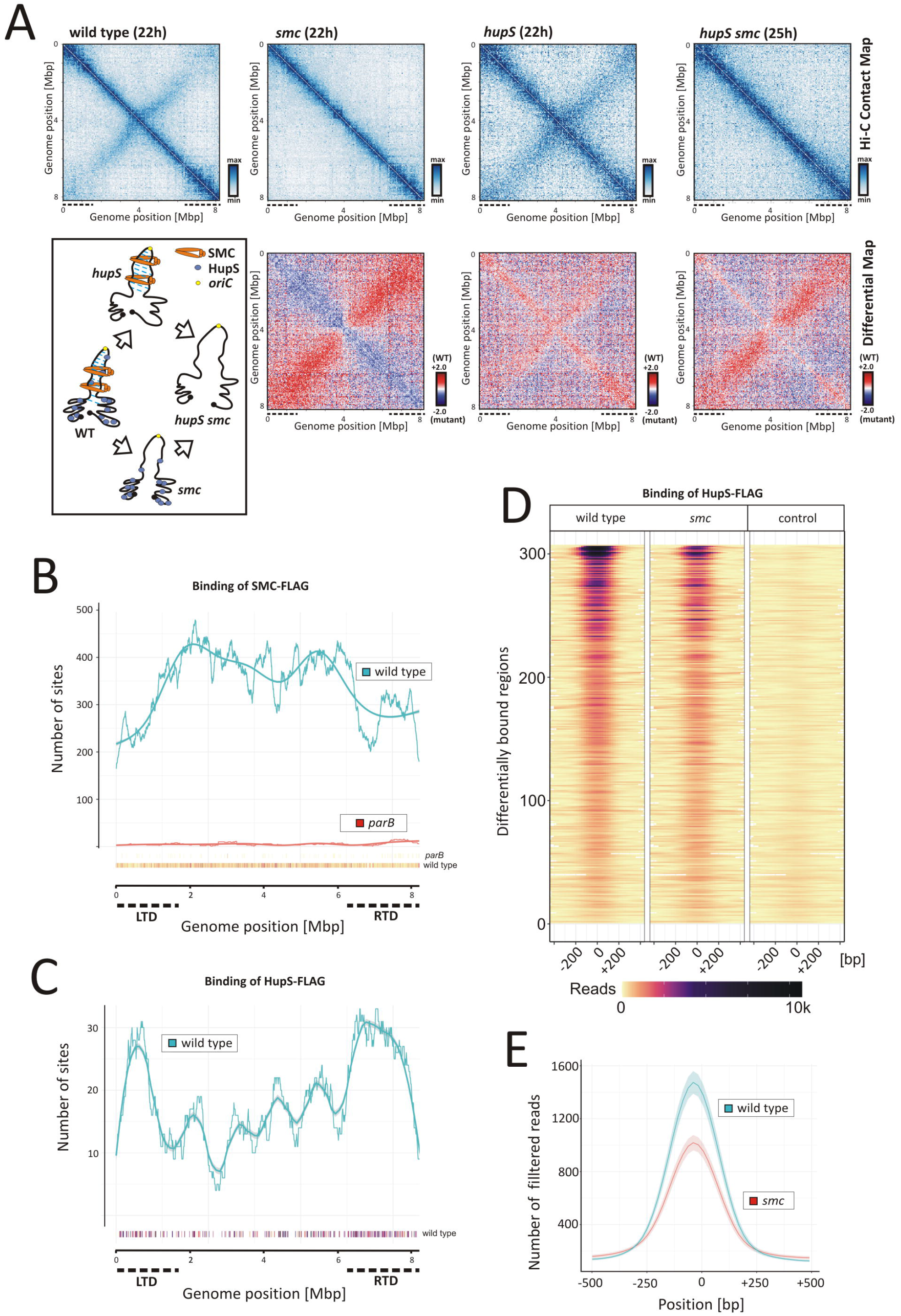
Influence of SMC and HupS on chromosome organization. (**A**) Top panels: the normalized Hi-C contact maps obtained for the wild type and *smc, hupS* and *smc hupS* double mutants (*ftsZ-ypet* derivatives, MD100, TM005, TM004, TM006, respectively) growing for 22 h or for 25 h (only in the case of the double *hupS smc* mutant) (in 5 ml cultures). Bottom panels: the corresponding differential contact maps in the logarithmic scale (log2) comparing the contact enrichment in the wild type strain (red) versus the mutant strains (blue) are shown below each Hi-C contact map. **(B)** The number of SMC-FLAG binding sites along the chromosome of the wild type (blue) (TM017 strain) or *parB* mutant background (red) (KP4F4) determined by ChIP-Seq analysis at the time of sporogenic cell division (14^th^ hour of 50 ml culture growth). The number of SMC-FLAG binding sites (250 bp) was determined by *normr* analysis averaged over 0.5 Mbp sliding window every 1000 bp with a loess model fit. Points below the plot show positions of individual sites coloured according to their FDR (false discovery rate) values (yellow less significant, dark purple more significant). (**C)** The number of HupS-FLAG binding sites along the chromosome in the wild type background (*hupS* deletion complemented with *hupS-FLAG* delivered *in trans*, TM015) determined by ChIP-Seq analysis at the time of sporogenic cell division (14^th^ hour of 50 ml culture growth). The number of HupS-FLAG binding sites was determined by differential analysis of the HupS-FLAG and wild type strains using *edgeR* and averaged over 0.5 Mbp sliding window every 1000 bp. Smooth line shows loess model fit. The points below the plot show positions of individual sites coloured according to their FDR values (yellow less significant, dark purple more significant). (**D**) Heatmaps showing the number of reads for all 307 regions bound by HupS-FLAG in the wild type background (TM015), *smc* mutant background (TM016) and negative control (the wild type strain). The reads were normalized by the glmQL model from the *edgeR* package. For each region, position 0 is the position with the maximum number of reads. Regions are sorted according to their logFC values. (**E**) Comparison of HupS-FLAG binding in the wild type (blue) and *smc* deletion background (red). The lines show mean values of reads (for 69 bp long regions) with 95% confidence intervals for all 307 sites differentially bound by HupS-FLAG protein. For all sites, 1000 bp long fragments were extracted from the chromosome centred around the best position as determined by *edgeR* analysis.

**Figure 4.**
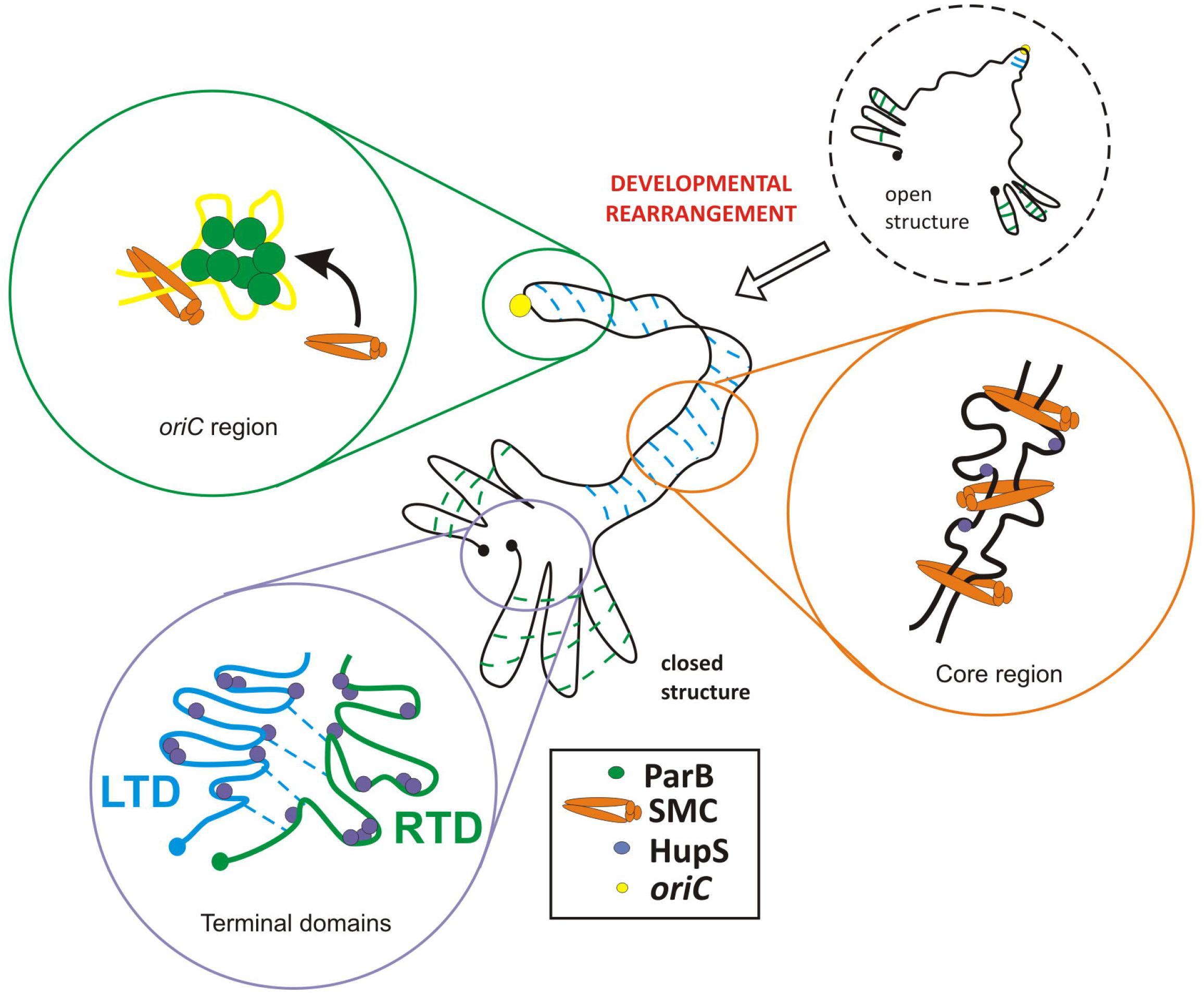
Model of spatial rearrangement of the *S. venezuelae* chromosome during sporogenic development showing the contribution of ParB (green), SMC (orange) and HupS (blue) to chromosomal arm alignment and organization of terminal domains (TLD and RTD). Blue lines indicated intrachromosomal contacts, *oriC* region is marked with yellow circle.

The Hi-C contact map generated for the *hupS* mutant at the time sporogenic cell division (at the 22^nd^ hour) revealed the presence of both diagonals. However, the identified local signals within both diagonals were more dispersed than in the wild type, and diagonals were visibly broadened. Moreover, along the primary diagonal, a less clear distinction between the core region and terminal domains could be observed. The effect of *hupS* deletion, though affecting whole chromosome compaction, was most prominent within terminal domains, especially within RTD (**Fig. 3A**).

To confirm HupS involvement in the organization of terminal domains and to test the cooperation between HupS and SMC, we analysed HupS-FLAG binding in the complemented *hupS* and double *hupS smc* deletion strains at the time of cell division. Notably, in contrast to SMC, HupS-FLAG showed low sequence binding preference, and we identified 307 regions differentially bound by HupS-FLAG with a mean width of 221 bp (**Fig. 3C, D** and **S8A**). The analysis of the number of HupS-FLAG binding sites exhibited a significant increase in the terminal domains, especially the right terminal domain, but HupS binding was also detected in the core region of the chromosome (**Fig. 3C**). The enhanced HupS binding at chromosomal termini is consistent with the lowered number of chromosomal contacts in RTD observed in the Hi-C maps obtained for the *hupS* mutant.

Further analysis of ChIP-seq data performed using two independent methods (*rGADEM* and *MEME*) identified the motif recognized by HupS-FLAG (**Fig. S8B**). Although the motif was observed to be uniformly widespread along the chromosome, and was not required for HupS binding, the analysis of probability showed its enrichment in HupS binding sites versus random regions and its preferred occurrence in the middle of the identified HupS-bound region (confirmed with *CentriMo*) (**Fig. S8B-E**). Additionally, *in vitro* studies showed that immobilized GST-HupS bound digested cosmid DNA with no sequence specificity (**Fig. S8F**). Markedly, ChIP-seq analysis of HupS-FLAG binding in the absence of SMC (*hupS smc* deletion complemented with *hupS*-FLAG) showed a lower number of reads for all HupS-FLAG binding sites in the absence of SMC, indicating that SMC enhanced the HupS-DNA interaction (**Fig. 3E**). This observation could not be explained by the low level of HupS-FLAG in the *smc* deletion background (**Fig. S9A**). We also confirmed earlier observations that the HupS-FLAG level during the pre-sporulation phase was lower than that observed during sporogenic cell division^42^, which corresponded with the marginal effect of *hupS* deletion on chromosome organization at the early pre-sporulation phase, as indicated by the Hi-C contact map for the *hupS* mutant at this time point (**Fig. S9B**).

Since ChIP-seq analysis indicated that the cooperation between HupS and SMC binding and double *hupS smc* deletion most severely affected chromosome compaction, we investigated how double deletion affects chromosomal arrangement. The Hi-C contact map generated for the double *hupS smc* mutant at the time of sporogenic cell division (at the 22^nd^ hour) showed the combination of effects observed independently for *hupS* and *smc* single mutations, namely, the disruption of arm alignment and the destabilization of local DNA contacts (**Fig. 3A**). Thus, while long-range interarm interactions are executed by SMC, short-range compaction is governed by HupS, and the effect of the elimination of both proteins is additive.

In summary, the Hi-C and ChIP-seq analyses of *smc* and *hupS* mutants showed their contribution to global chromosome compaction. While SMC binds in the chromosomal core and is responsible for the juxtaposition of arms, HupS binds in terminal domains and organizes them. The cooperation between both proteins is manifested by SMC enhancing HupS binding to DNA. Elimination of both proteins disturbs global chromosome organization.

## DISCUSSION

In this study, we provide a complex picture of the dynamic rearrangement of linear bacterial chromosome. The Hi-C mapping of the ∼8.2 Mbp *S. venezuelae* chromosome at the time of sporogenic cell division revealed arm juxtaposition, which extends approximately 4 Mbp around the *oriC* region. The interarm contacts are diminished at the independently folded 1.5 and 2 Mbp C-terminal domains LTD and RTD, respectively. Further analysis of chromosomal contacts shows spatial proximity of both chromosomal termini, which is consistent with the findings of previous cytological studies^45^. Folding of the *Streptomyces* linear chromosome into core and terminal domains was also independently demonstrated for the *S. ambofaciens* (see Lioy *et al*., submitted back to back).

Our Hi-C data disclosed profound rearrangement of the chromosomal conformation that occurs during *S. venezuelae* sporulation. At the end of vegetative growth, at the entrance to the sporulation phase contacts between the chromosomal arms are scarce. After this time point, the chromosomal arms become increasingly aligned, reaching the maximum at the time of initiation of sporogenic cell division (Z-ring formation) which is also the time point when chromosome segregation is initiated^39^. Thus, during *Streptomyces* differentiation, the changes in hyphal morphology, gene expression, chromosome replication and mode of cell division^14,34^ are accompanied by the rearrangement of chromosomes from their extended conformation during vegetative growth to almost fully folded and compacted structures.

Our data showed that in *S. venezuelae*, both ParB and SMC are essential for chromosomal arm juxtaposition, reinforcing the ParB-dependent SMC loading onto the chromosome. Importantly, in S. *venezuelae*, SMC binding is enriched within the core region of the chromosome, reflecting the extension of interarm contacts. SMC was previously shown to be involved in chromosome compaction during *S. coelicolor* sporulation^40,71^. Interestingly, *parB* deletion was also observed to decrease chromosome condensation at the time of cell division^39^. This observation may be explained by the lack of SMC loading in absence of ParB, which was also reported for the other model bacteria^4,6,7,9^. The increase in long-range chromosomal contacts induced by SMC in the chromosomes of *S. venezuelae* and other bacteria corroborates proposed model of SMC action which zips DNA regions when translocating from the loading site^4^.

While the increase of arm alignment during sporulation cannot be explained by elevation of SMC level during sporulation, the *parAB* genes were earlier shown to be transcriptionally induced in *Streptomyces* sporogenic hyphae^65,75^. ParB binds to *oriC* regions of all chromosomes throughout the whole life cycle but complexes formed during vegetative growth and at the early stages of sporogenic development are not as massive as regularly spaced segrosomes that accompany sporogenic-associated cell division^70^. Since interarm contacts occur during cell division, characteristic spatial architecture of sporogenic segrosomes is implied to be a prerequisite for SMC loading. The significance of the ParB complex architecture for SMC loading has been established by studies of ParB mutants with impaired spreading or bridging in *B. subtilis* and in *C. glutamicum*^7,27^. Therefore, our observations suggest that during *Streptomyces* differentiation ParB complexes spatially rearrange to increase their capability of SMC loading and initiating chromosomal arm juxtaposition.

What factors could induce the reorganization of ParB complexes during *Streptomyces* sporulation? The recently shown ParB ability to bind and hydrolyse CTP that affects formation of segrosomes and appears to be conserved among ParB homologs^76–78^ while not yet confirmed in *Streptomyces* may be involved in the rearrangement of ParB complexes in their sporogenic hyphae. On the other hand, the formation of ParB complexes was also shown to be associated with sporulation-dependent deacetylation of ParB in *S. coelicolor*^82^. The factor contributing to segrosomes rearrangement may be the increased transcriptional activity of the *parAB* operon. ParA accumulates in sporogenic cells, promoting segrosome formation and their regular distribution^39,70^. In other bacteria (*B. subtilis, V. cholerae, and M. smegmatis*) the elimination of ParA also decreased ParB binding^79–81^. Indeed, our Hi-C mapping showed that in the absence of ParA, the interarm interactions are somewhat diminished, reinforcing the role played by ParA in executing ParB architecture. Finally, since transcriptional activity was proven to have a considerable impact on chromosomal arm alignment^5,83^ (see also Lioy *et al*., submitted back to back), sporulation-associated changes in transcription within the chromosomal core may account for the development of interarm contacts.

We established that HupS, unique actinobacterial sporulation-specific HU homologue^42^, although promotes contacts all over the chromosome, is particularly involved in the organization of the terminal domains. HupS exhibited binding sequence preference *in vivo*, but no sequence specificity when binding linear DNA fragments. Thus, we infer that specific DNA structural features may account for enhanced HupS binding in particular regions. Lowered HupS binding in the absence of SMC could be explained by preferential binding by HupS to structures such as loops, generated by SMC. This notion corroborates earlier suggestions that HU homologues could stabilize DNA plectonemic loops, increasing short-range contacts^6^. *E. coli* studies also suggested cooperation between condensin MukB and HU homologues in maintaining long-range chromosomal contacts^10^. In fact, in *E. coli*, condensin activity was suggested to be increased by DNA structures induced by HU protein^10^. Previous analysis of *S. coelicolor* showed that HupS deletion resulted in chromosome decompaction in sporogenic hyphae^42^. The Hi-C matrixes established that both proteins, SMC and HupS contribute to chromosome compaction during sporulation, and the double *hupS smc* deletion has the strongest effect on chromosomal conformation, affecting both arm juxtaposition and short-range chromosomal contacts.

Although elimination of SMC and HupS disturbed the *S. venezuelae* chromosome structure, the phenotype of the double mutant was still rather mild and manifested predominantly in increased nucleoid volume and formation of elongated spores. While the mild phenotype of the *smc* mutant was also described in *C. glutamicum*, in most bacteria (e.g., *B. subtilis. C. crescentus*, and *S. aureus*), *smc* deletion resulted in either chromosome decompaction and/or segregation defects and often led to the formation of elongated cells^7,84,85^. Interestingly, condensins were suggested to be particularly critical for chromosome segregation in bacteria lacking the complete *parABS* system (such as *E. coli*)^86–88^. The observed in *S. venezuelae smc* mutants aberrations of spore size correlate with irregular placement and lowered stability of Z-rings, suggesting that decreased chromosome compaction affects Z-ring formation and/or stability.

In summary, we established maps of chromosomal contacts for linear bacterial chromosomes. The *S. venezuelae* chromosome conformation undergoes extensive rearrangement during sporulation. Similar chromosomal rearrangement was observed during the transition from the vegetative to stationary phase of growth in *S. ambofaciens* (Lioy *et al*. submitted back to back). DNA organizing proteins HupS and SMC contribute to chromosome compaction, and they also impact cell division. Although HU and SMC roles have been established for numerous bacteria, in *Streptomyces*, their function is seemingly adjusted to meet the requirements of compacting the large linear chromosomes. Due to the unique *Streptomyces* differentiation, in which chromosome replication and segregation are spatiotemporally separated, the transition between the particular stages of life cycle requires the chromosomal rearrangement similarly profound as rearrangement of chromosomes during eukaryotic cell cycle.

## Supporting information

Supplementary_Informations

## DATA DEPOSITION

Raw data are available at ArrayExpress (EMBL-EBI) with accession numbers: E-MTAB-9810 (Hi-C data), E-MTAB-9821 (ChIP-Seq data, HupS-FLAG immunoprecipitation), and E-MTAB-9822 (ChIP-Seq data, SMC-FLAG immunoprecipitation).

## ACKNOWLEDGEMENTS

We are grateful to Susan Schlimpert for sharing pSS170 and pSS172*hupA-mcherry*. We thank the Bioimaging facility of the John Innes Centre, supported by a core capability grant from BBSRC, for use of their resources. T.B.K.L. acknowledges the Royal Society University Research Fellowship (UF140053). This work was funded by the Polish National Science Centre: HARMONIA grant 2016/22/NZ1/00122 (to M.J.S.), OPUS grant 2018/31/B/NZ1/00614 (to D.J.).

