## Supplementary_Figures for "Spatial rearrangement of the *Streptomyces venezuelae* linear chromosome during sporogenic development"

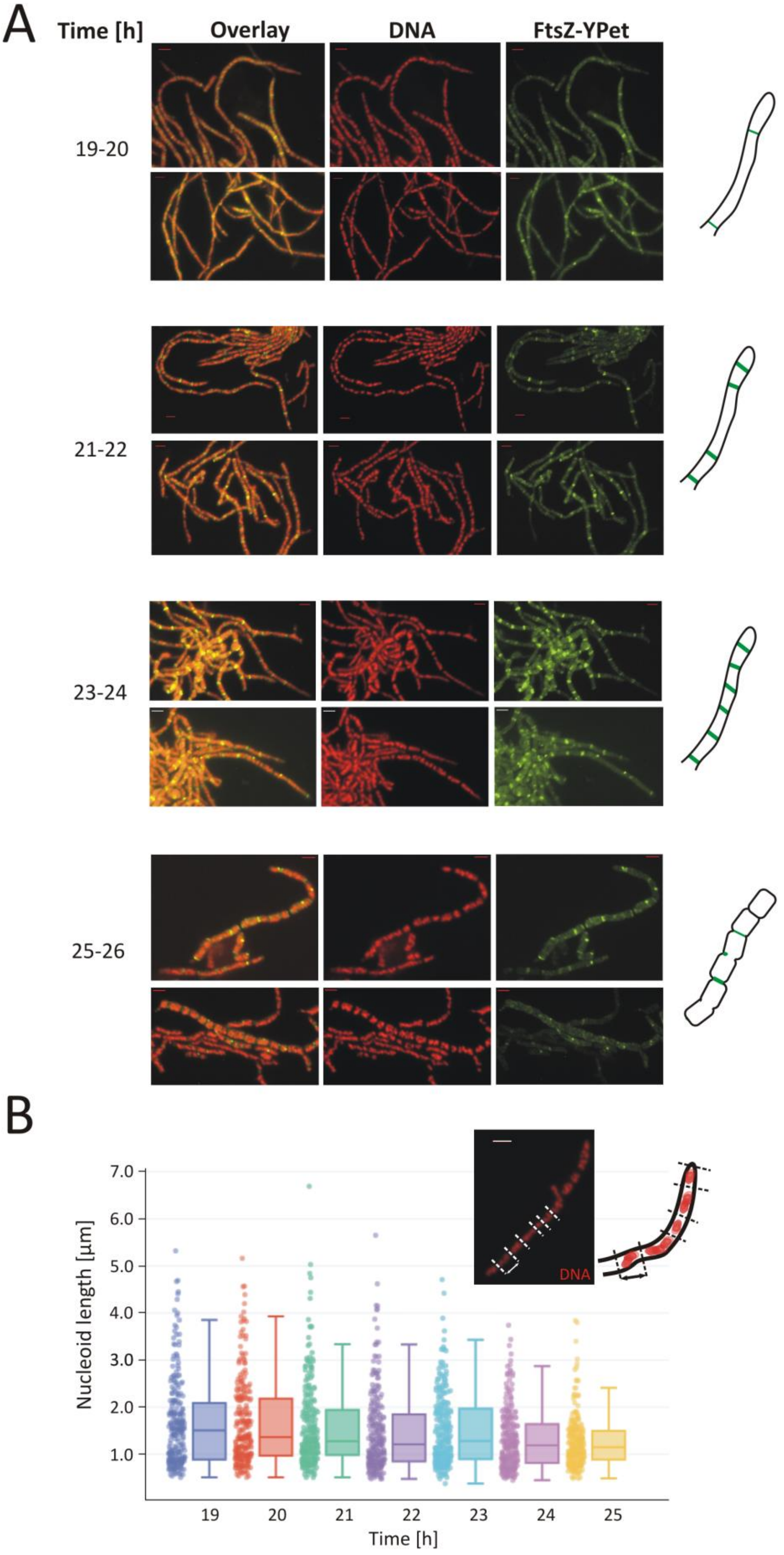
A
Time [h]
Overlay
DNA
FtsZ-YPet
19-20
21-22
23-24
25-26
B
Nucleoid length [µm]
7.0
6.0
5.0
4.0
3.0
2.0
1.0
19
20
21
22
23
24
25
Time [h]
DNA

Figure S1

**Figure S1. Formation of Z-ring ladders and nucleoid compaction during *S. venezuelae* sporulation**. **(A)** Fluorescence microscopy images of the wild type strain (*ftsZ-ypet* derivative, MD100) hyphae fixed at different time points of culture (5 ml culture). The images show the visualization of nucleoids (DNA stained with 7-AAD) and the FtsZ-YPet signal. The schematic drawings on the right side show the hyphae at particular time points of *S. venezuelae* sporogenic development. (**B**) Analysis of nucleoid compaction at the different time points of *S. venezuelae* growth. The nucleoid compaction was quantified as the distance (marked with a black arrow) between the centres of two DNA-free zones. The inset shows a representative picture showing an example of the data collection. Scale bar - 2 µm.

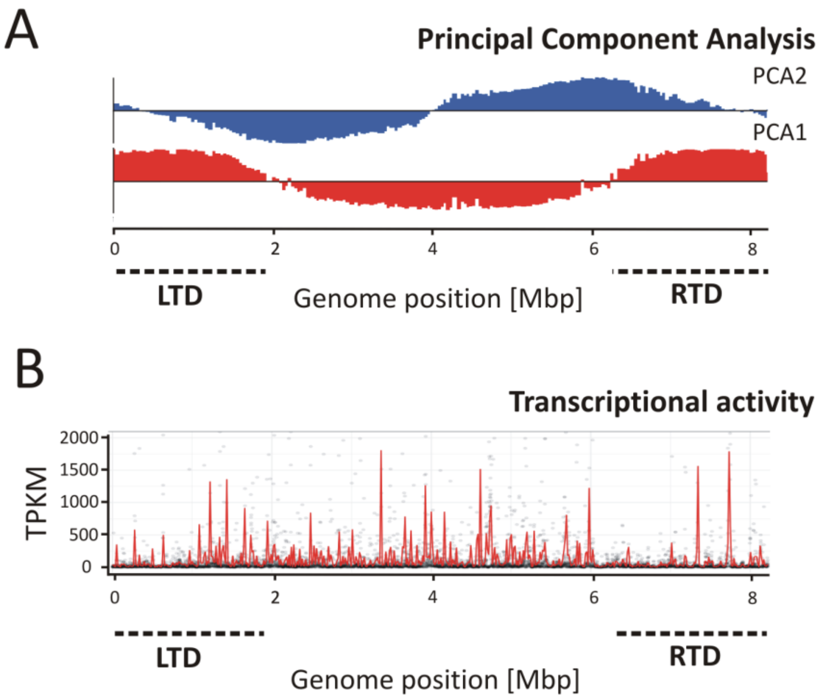


Figure S2

**Figure S2. Identification of distinct regions within the *S. venezuelae* chromosome.** (**A**) Principal component analysis (PCA1 - red, PCA2- blue) of the corrected and normalized Hi-C matrix in from the wild type (*ftsZ-ypet* derivative, MD100) strain growing for 22 h (5 ml cultures). The positions of the left (LTD) and right (RTD) terminal domains are marked with black dotted lines. (**B**) The transcriptional activity in the *S. venezuelae* wild type strain measured as the number of transcripts per kilobase million (TPKM). The red line shows the average TPKM calculated in a 15-kbp sliding window. The analysis was based upon processed data from project E-MTAB-7457[63].

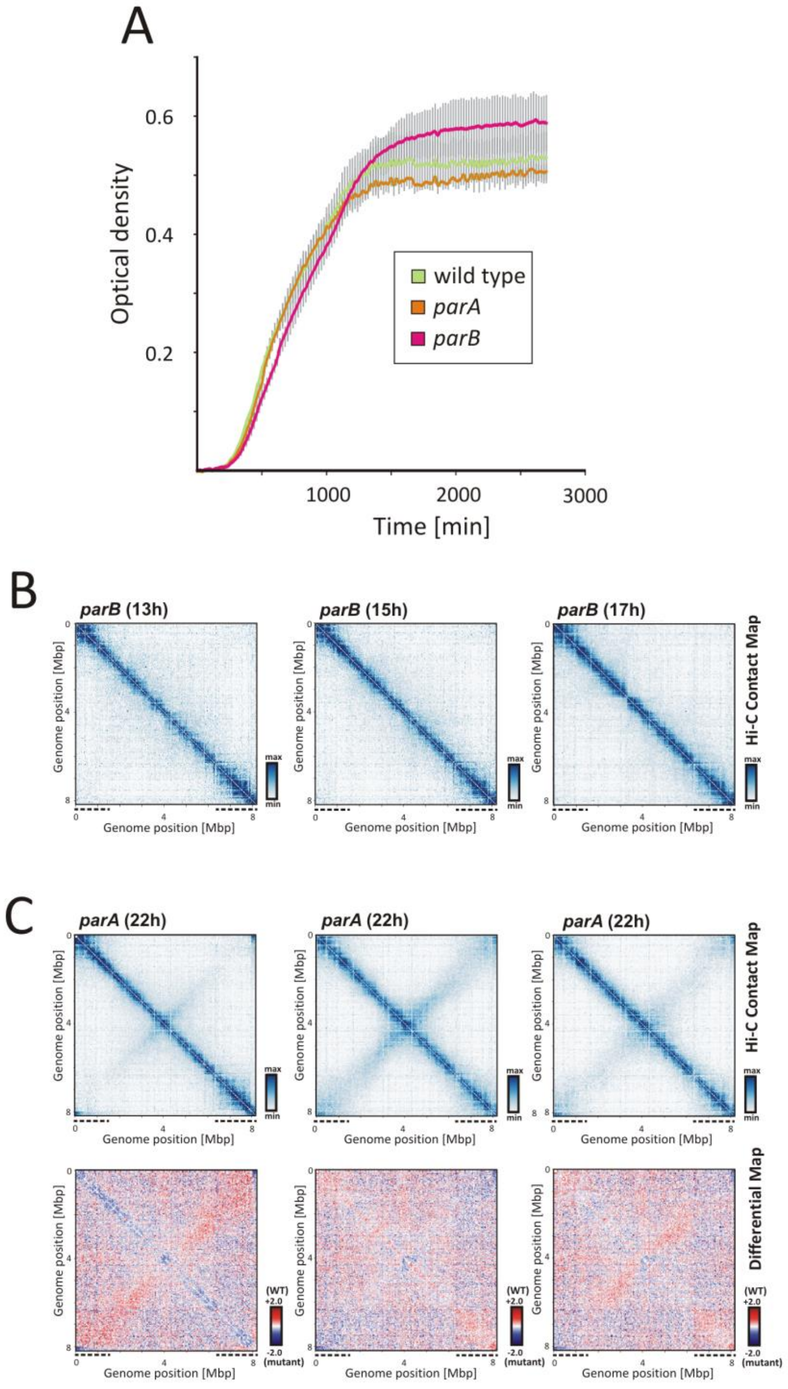


Figure S3

**Figure S3. Growth and chromosome organization of *parA* and *parB* mutants. (A)** The growth of the wild type strain (green) as well as *parA* (orange) and *parB* (pink) in MYM medium quantified using a Bioscreen C instrument. All analysed strains are *ftsZ-ypet* derivatives (MD100, MD011 and MD021) **(B)** The normalized Hi-C contact maps obtained for the *parB* mutant (*ftsZ-ypet* derivative, MD021) after 13, 15, and 17 h of growth (5 ml culture). **(C)** Top panel: the normalized Hi-C contact maps obtained for three biological repeats of the *parA* mutant (*ftsZ-ypet* derivative, MD011) growing for 22 h (5 ml culture). Bottom panel: the differential contact maps in the logarithmic scale (log2) comparing the contact enrichment in the wild type strain (red) versus the *parA* mutant (blue) are shown below each replicate.

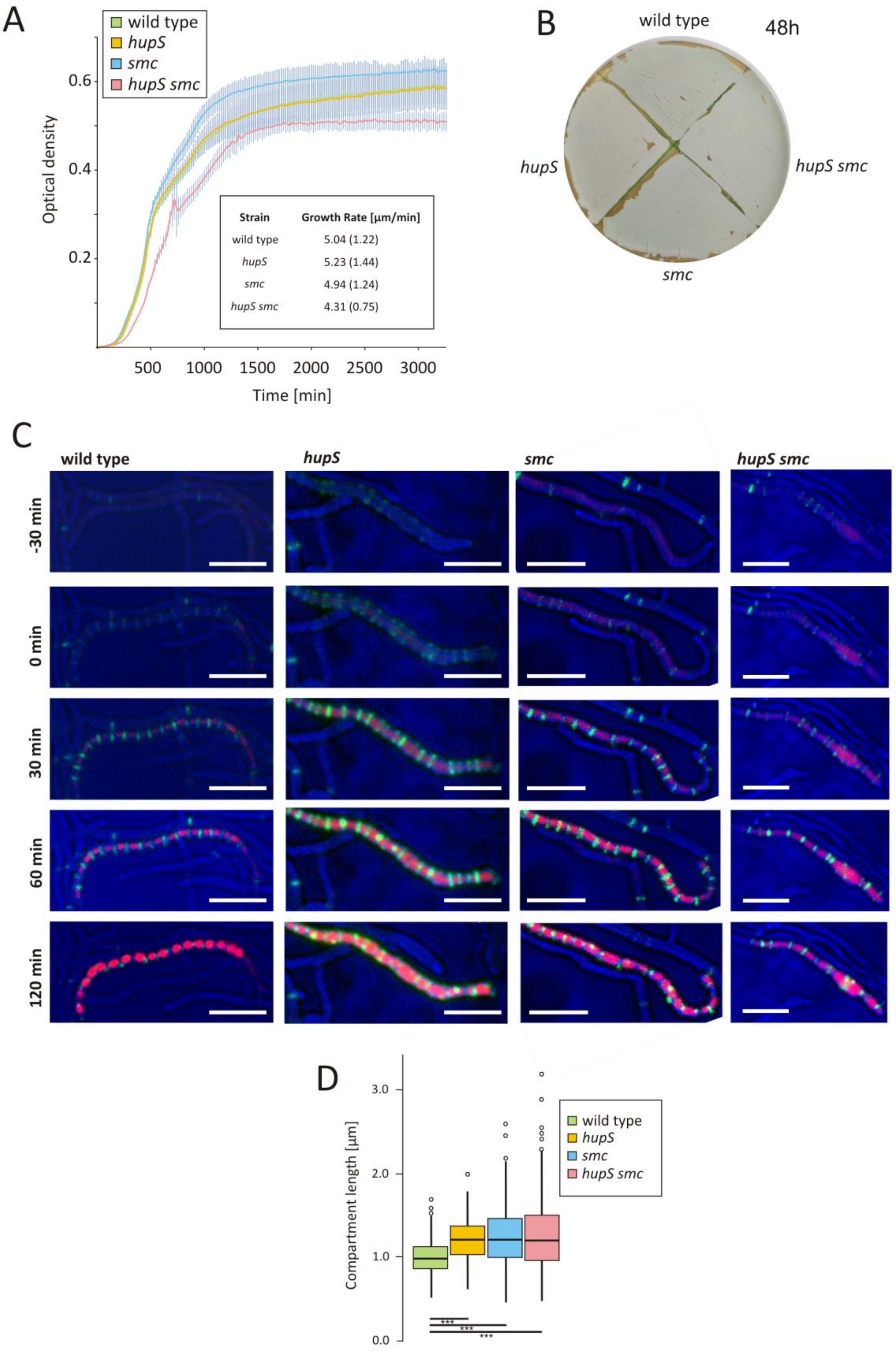
A
wild type
hupS
smc
hupS smc
Optical density
0.2
0.4
0.6
500
1000
1500
2000
2500
3000
Time [min]
Strain
Growth Rate [µm/min]
wild type
5.04 (1.22)
hupS
5.23 (1.44)
smc
4.94 (1.24)
hupS smc
4.31 (0.75)
B
wild type
48h
hupS
hupS smc
smc
C
wild type
hupS
smc
hupS smc
-30 min
0 min
30 min
60 min
120 min
D
Compartment length [µm]
0.0
1.0
2.0
3.0
wild type
hupS
smc
hupS smc
***
***
***

Figure S4

**Figure S4. Detailed analysis of the phenotypic effects of *smc*, *hupS* and double *smc hupS* deletion. (A)** The growth of the wild type (light green), *hupS* (yellow), *smc* (blue) and *hupS smc* (pink) double mutant (*ftsZ-ypet* derivatives, MD100, TM005, TM004, TM006) in MYM medium quantified using a Bioscreen C instrument. The inset shows the average growth rate [µm/min] of a germ tube observed in the time-lapse experiment. Standard deviations are shown in brackets. **(B)** The growth of the wild type as well as *hupS*, *smc* and *hupS smc* double mutant analysed after 48 h of growth on solid MYM medium (AKO200, TM010, TM003, respectively). **(C)** Time-lapse observations of the wild type as well as *hupS*, *smc* and *hupS smc* double mutant (*ftsZ-ypet*, *hupA-mCherry* derivatives, TM011, TM013, TM012 and TM014, respectively). Nucleoid condensation was visualized using mCherry-HupA fusion, whereas Z-rings were visualized using FtsZ-YPet fusion. The representative images show the hyphae 30 min before and up to 130 min after the growth arrest of the sporogenic compartment (time = 0 min). Scale bar: 5 µm. **(D)** Box plot analysis of the compartment length distribution. The distance between two Z-rings was measured during the sporulation of the wild type as well as *hupS*, *smc* and *hupS smc* double mutant (*ftsZ-ypet* derivatives, MD100, TM005, TM004, TM006, respectively).

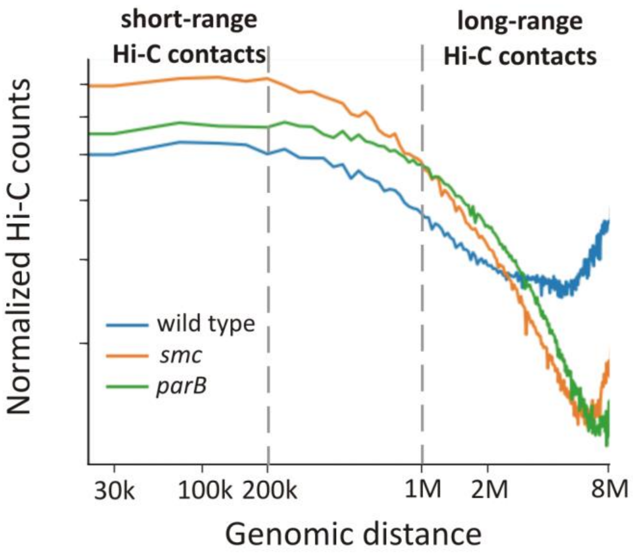


### Figure S5

**Figure S5. Involvement of SMC and ParB in short- and long-range DNA contacts.** (**A**) The normalized Hi-C contacts calculated using the *hiCPlotDistVsCounts* package (Galaxy Version 3.4.3.0) for the wild type strain (blue), *smc* (orange) and *parB* (green) mutants (*ftsZ-ypet* derivatives, MD100, TM004 and MD021, respectively). The X-axis shows the genomic position on a logarithmic scale. The short-range (below 200 kbp) and long-range Hi-C contacts (above 1 Mbp) are marked with dotted lines.

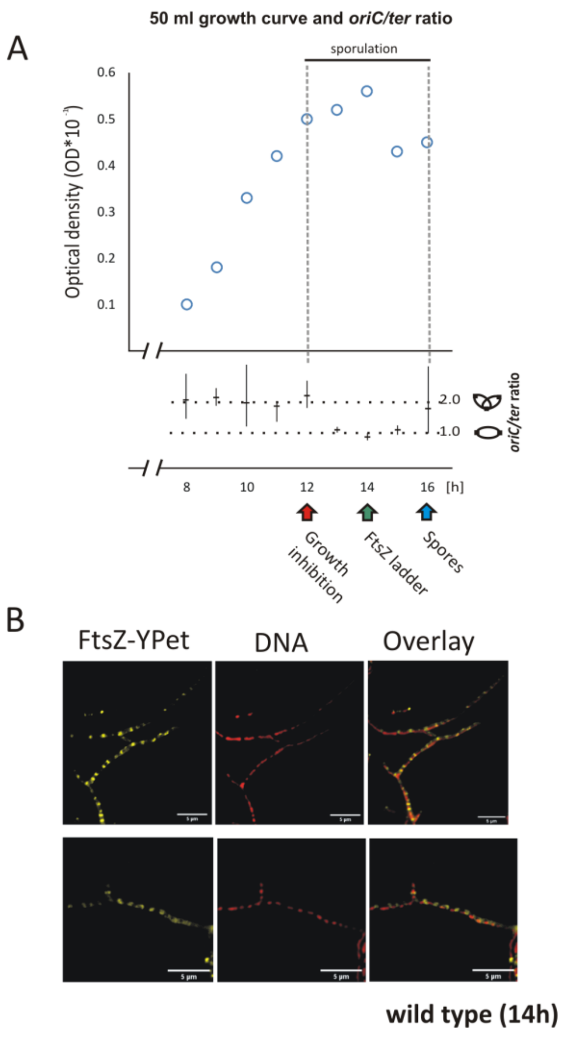


### Figure S6

**Figure S6. Standardization of the *S. venezuelae* “ChIP-seq culture”. (A)** The growth curve under ChIP-seq culture conditions (50 ml cultures). The critical time points of sporogenic development are marked with arrows: growth arrest (red), the appearance of FtsZ ladders (green) and the formation of spore chains (blue). The relative *oriC*/*ter* ratio is shown below the plot, with X axes corresponding to the main plot X axis. The *oriC/ter* ratio at the 16$^{th}$ hour of growth was set as 1.0. **(B)** Fluorescence microscopy images of the wild type (*ftsZ-ypet* derivative MD100) hyphae fixed at the 14$^{th}$ hour of growth (50 ml culture). The images show the visualization of nucleoids (DNA stained with 7AAD) and the FtsZ-YPet signal. Scale bar: 5 µm.

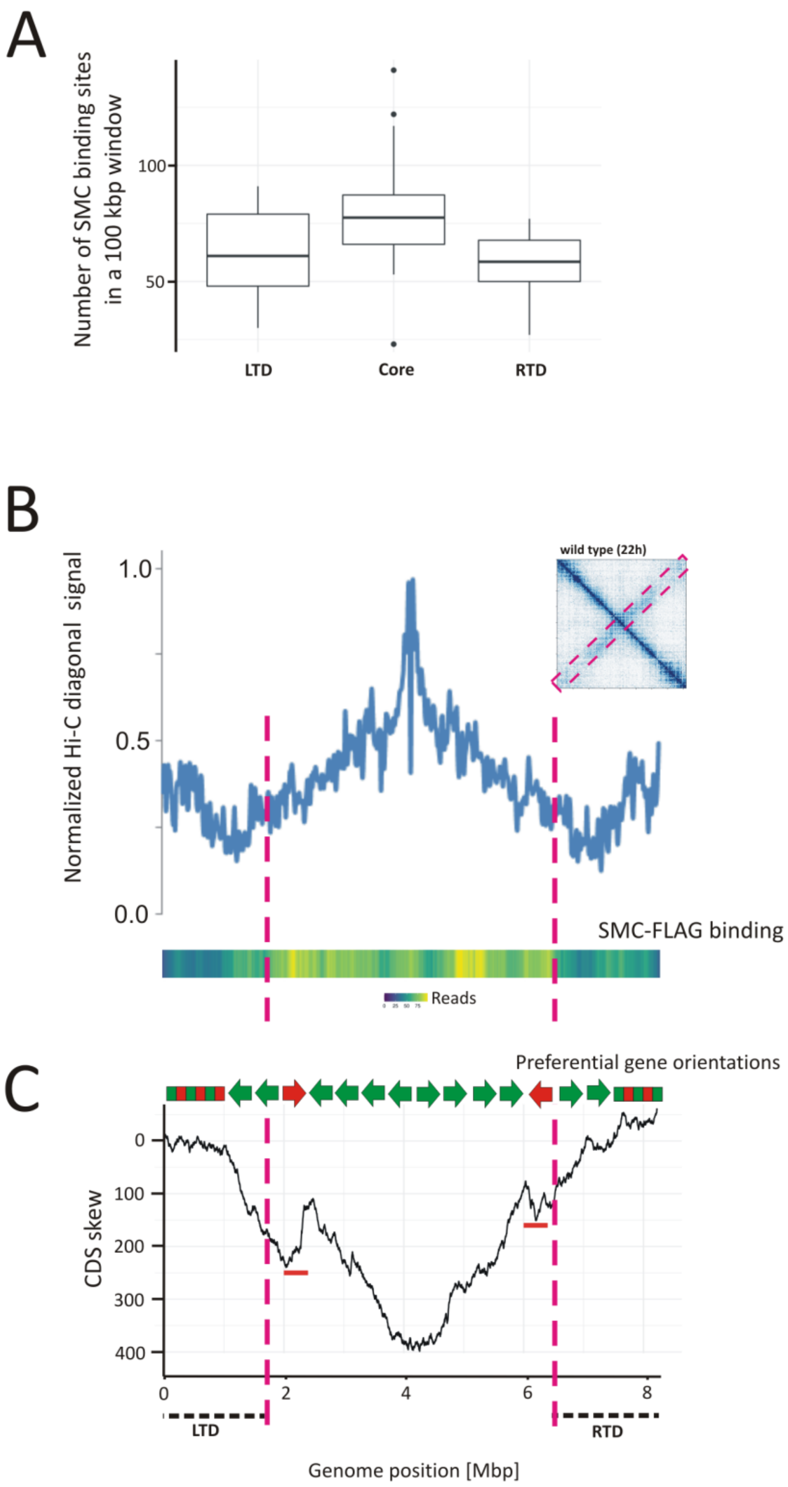
A
Number of SMC binding sites in a 100 kbp window
100
50
LTD
Core
RTD
B
Normalized Hi-C diagonal signal
1.0
0.5
0.0
wild type (22h)
SMC-FLAG binding
Reads
C
Preferential gene orientations
CDS skew
0
100
200
300
400
0
2
4
6
8
LTD
RTD
Genome position [Mbp]

Figure S7

**Figure S7. Overlapping SMC-FLAG binding sites with the Hi-C diagonal signal. (A)** Boxplot analysis of the number of SMC-bound regions (250 bp) counted in 0.1 Mbp window for core (1.9-6.3 Mbp), LTD (<1.9 Mbp) and RTD (>6.3 Mbp) domains. **(B)** The average value of the signal along the secondary diagonal axis was calculated using Fiji software based on the Hi-C contact map obtained for the wild type strain (*ftsZ-ypet* derivative, MD100), as shown in the insert at the 22nd hour of growth, and compared with the heat map of identified SMC-FLAG binding sites in the wild type background. The heat scale corresponds to the number of binding sites in the 0.5 Mbp sliding window every 1000 bp. The positions of the LTD and RTD domains are marked with black dotted lines. The pink dotted lines show the edges of LTD and RTD for easier comparison with Hi-C and ChIP-Seq data. **(C)** Cumulative plot of gene orientation bias (CDS skew) along *S. venezuelae* chromosome. The curve moves up one unit for each gene encoded on the forward strand and one unit down for each gene on the reverse strand. Regions of the chromosome containing genes located in the same (green arrow) or opposite (red arrow) direction to the preferential gene orientation are marked above. The terminal regions without preferences in gene orientations were marked with red-green rectangles.

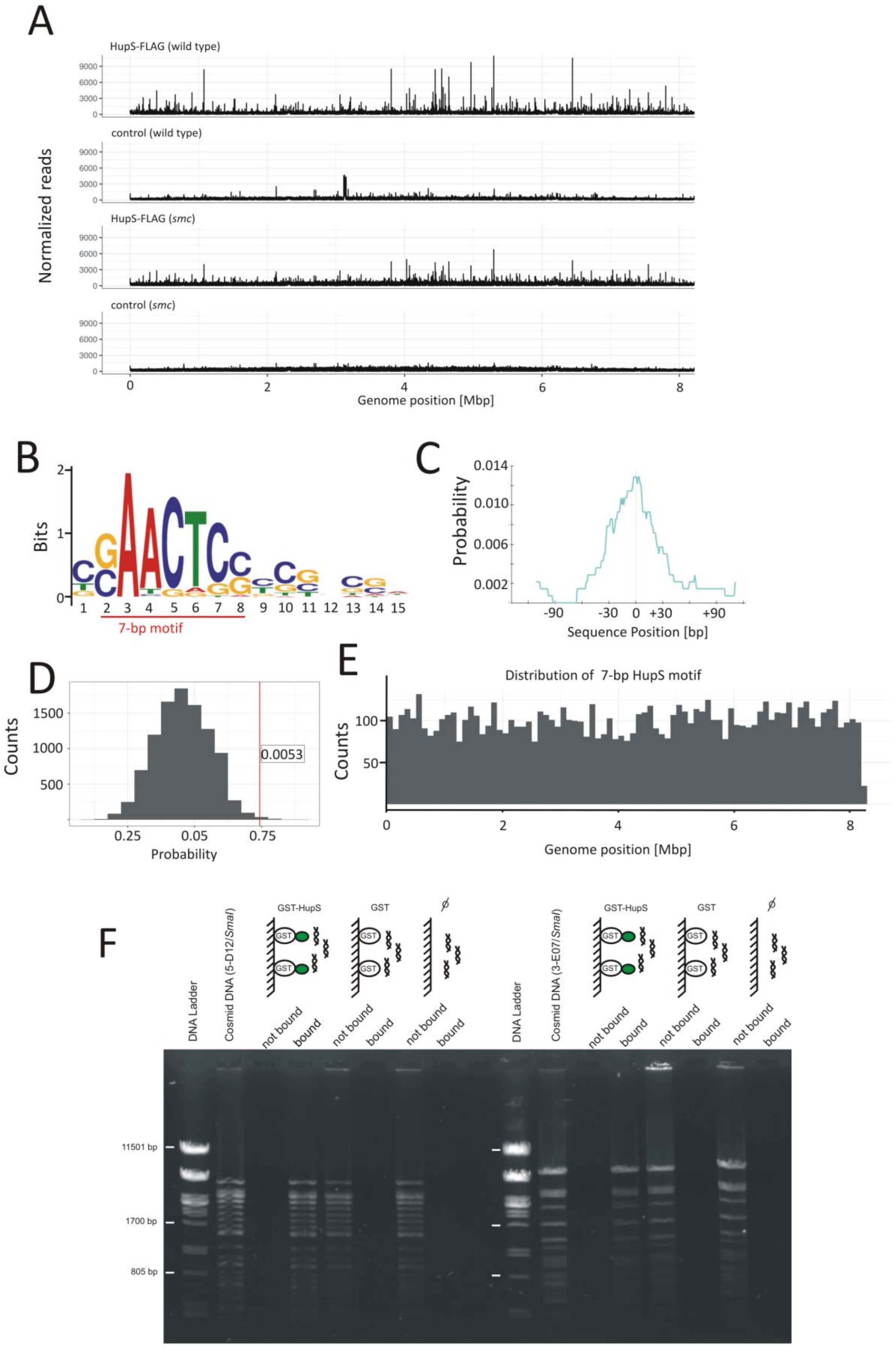
A
HupS-FLAG (wild type)
control (wild type)
HupS-FLAG (smc)
control (smc)
Normalized reads
Genome position [Mbp]
B
Bits
7-bp motif
C
Probability
Sequence Position [bp]
D
Counts
0.0053
Probability
E
Distribution of 7-bp HupS motif
Counts
Genome position [Mbp]
F
DNA Ladder
Cosmid DNA (5-D12/SmaI)
GST-HupS
GST
not bound
bound
Cosmid DNA (3-E07/SmaI)
11501 bp
1700 bp
805 bp

Figure S8

**Figure S8. Analysis of HupS-FLAG binding. (A)** The normalized ChIP-Seq reads for HupS-FLAG in the wild type background (TM015) and in the *smc* deletion background (TM016). Each experiment was supplemented with the control showing read distribution in the negative control strains of the wild type and *hupS* mutant (lacking *hupS-flag* gene). (**B**) Logo of the 15 bp sequence found by the *MEME suite* to be significantly enriched in HupS ChIP-identified regions, with the most conserved 7 bp motif indicated by the red line. (**C**) Local enrichment of probable HupS motifs in the centre of ChIP-identified regions. Plot produced by *CentriMo* software. (**D**) Permutation test showing enrichment of consensus sequences identified by the *MEME suite* in HupS ChIP-identified regions (red line, with calculated p-value shown) versus the random sequences (histogram). (**E**) Distribution of 7-bp motifs identified by the *MEME suite* across the entire S*. venezuelae* chromosome, counted in 0.1 Mbp windows. **(F)** Analysis of DNA binding by GST-HupS recombinant protein *in vitro*. The lysates from *E. coli* BL21 pLys overexpressing GST-HupS recombinant protein (or GST protein in the control experiment) were incubated with Glutathione Sepharose resin (GE Healthcare), washed to remove unbound proteins and subsequently incubated with cosmids (5-D12, containing multiple HupS consensus sequences, and 3-E07 lacking these sequences) digested earlier with SmaI restriction enzyme. The unbound and bound (eluted with 2 M NaCl) fractions were analysed in a 1% agarose gel and visualized against DNA ladder (phage lambda DNA digested with PstI restriction enzyme) with ethidium bromide staining. As an additional control, Glutathione Sepharose resin was not incubated with *E. coli* cell lysates but only with digested cosmids.

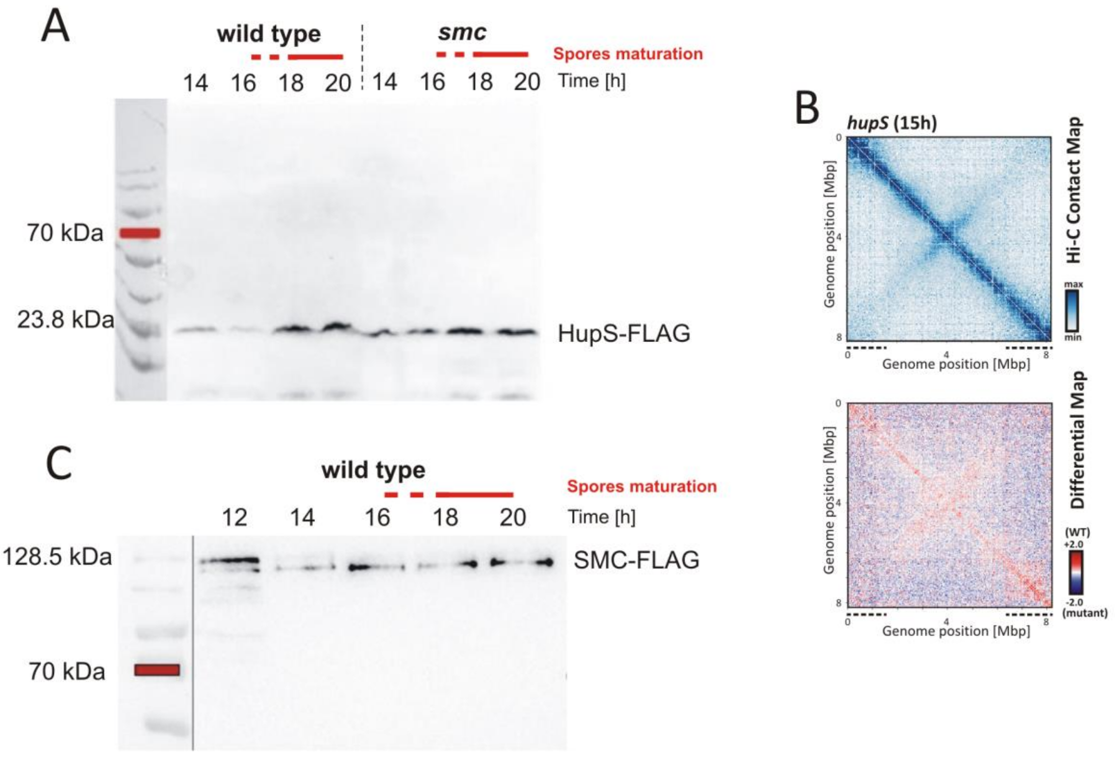


Figure S9

**Figure S9. HupS and SMC levels during sporulation. (A)** Western blot detection of HupS-FLAG using anti-FLAG monoclonal antibody in the wild type background (TM015) and *smc* mutant (TM016) (growing for 14-20 h in MYM liquid (5 ml culture). The sporulation phase was marked with the red line. **(B)** The normalized Hi-C contact maps obtained for the wild type and *hupS* mutant growing for 15 h (5 ml culture). The differential Hi-C map in the logarithmic scale (log2) comparing the contact enrichment in the wild type strain (red) versus the mutant strain (blue) is shown below. **(C)** Western blot detection of FLAG-SMC using anti-FLAG monoclonal antibody in the wild type background (TM017) growing for 12-20 h in MYM liquid (5 ml culture). The sporulation phase was marked with the red line.

**Supplementary Movies 1-4.** Time-lapse observations of the wild type (Movie 1) as well as *hupS* (Movie 2), *smc* (Movie 3) and *hupS smc* double mutant (Movie 4) ((*ftsZ-ypet*, *hupA-mCherry* derivatives, TM011, TM013, TM012 and TM014, respectively). Nucleoid condensation was visualized using mCherry-HupA fusion, whereas Z-rings were visualized using FtsZ-YPet fusion. The images were taken every 10 minutes. Scale bar: 5 µm.
