## Supplementary_Informations for "Spatial rearrangement of the *Streptomyces venezuelae* linear chromosome during sporogenic development"

**Supplementary Information**

**Table S1** ***Streptomyces venezuelae*** **strains used in the study**

| **Short name/strain number** | **Relevant genotype and characteristics** | **source** |
|---|---|---|
| wt / - | Wild type S. *venezuelae*<br><br>NRRL B-65442 (number in NRRL culture collection , genome *NZ_CP018074.1*) | Kind gift from prof. Mark Buttner, John Innes Centre, Norwich, UK [1] |
| *ftsZ-ypet* / **MD100** | *attBΦC31*:: pKF351 *ftsZ-ypet -apra* (Apr$^R$) | [2] |
| *parB* (*ftsZ-ypet*) / **MD021** | *ΔparB::apra, attBΦC31::* pKF351 *ftsZ-ypet hyg* (Hyg$^R$ , Apr$^R$) | [2] |
| *parA* (*ftsZ-ypet*) / **MD011** | *ΔparA, attBΦC31:: pKF351 ftsZ-ypet hyg* (Hyg$^R$) | [2] |
| - / **TM001** | *Δsmc::apra* (Apr$^R$), | This study |
| *smc* / **TM010** | *Δsmc::scar* (modified TM001) | This study |
| *hupS* / **AKO200** | *ΔhupS*::*apra* (Apr$^R$) | This study |
| *hupS smc* / **TM003** | *Δsmc::scar, ΔhupS*::*apra* (modified TM010) (Apr$^R$) | This study |
| *smc* (*ftsZ-ypet*) / **TM004** | Δ*smc::scar, attBΦC31*:: pKF351 *ftsZ-ypet apra* (modified TM010) (Apr$^R$) | This study |
| *hupS* (*ftsZ-ypet*) / **TM005** | Δ*hupS::apra, attBΦC31*:: pKF351 *ftsZ-ypet hyg* (modified AKO200) (Hyg$^R$) | This study |
| *hupS smc* (*ftsZ-ypet*) / **TM006** | Δ*smc::scar*, Δ*hupS::apra*, *attBΦC31*:: pKF351 *ftsZ-ypet-hyg* (modified TM003) (Hyg$^R$ , Apr$^R$) | This study |
| wt (*ftsZ-ypet hupA-mcherry*) / **TM011** | *attBΦC31*:: pKF351 *ftsZ-ypet apra, ttBφBT1*::pSS172*hupA-mchery* (modified MD100) (Apr$^R$, Hyg$^R$) | This study |
| *smc* (*ftsZ-ypet hupA -mcherry*) / **TM012** | Δ*smc::scar*, *attBΦC31*:: pKF351 *ftsZ-ypet apra*, *attBφBT1*::pSS172*hupA -mcherry,* (modified TM010) (Apr$^R$, Hyg$^R$) | This study |
| *hupS* (*ftsZ-ypet hupA-mcherry*) / **TM013** | Δ*hupS::apra*, *attBΦC31*:: pKF351- *ftsZ-ypet* spec, *attBφBT1*::pSS172*hupA-mcherry* (modified AKO200) (Apr$^R$, Hyg$^R$, Spec$^R$) | This study |
| *hupS smc* (*ftsZ-ypet hupA-mcherry*) / **TM014** | Δ*smc::scar*, *attBΦC31*:: pKF351 *ftsZ-ypet spec, attBφBT1*::pSS172 *hupA-mcherry* (modified TM003) (Apr$^R$, Hyg$^R$, Spec$^R$) | This study |
| *hupS-FLAG* / **TM015** | Δ*hupS::apra, attBφBT1*::pSS172*hupS-FLAG* (modified AKO200)(Apr$^R$, Hyg$^R$) | This study |
| *hupS-FLAG smc* / **TM016** | Δ*hupS::apra*, Δ*smc::scar, attBφBT1*::pSS172 *hupS-FLAG* (modified TM003) (Hyg$^R$, Apr$^R$) | This study |
| *FLAG-smc* / **TM017** | *smc* :: *FLAG-smc* | This study |
| *hupS FLAG-smc* / **TM018** | *smc* :: *FLAG-smc,* Δ*hupS::apra* (modified TM017) (Apr$^R$) | This study |
| *parB FLAG-smc* / **KP4F4** | *smc* :: *FLAG-smc,* Δ*parB*::*apra,* modified TM017) (Apr$^R$) | This study |

| | | |
|---|---|---|
| - / **TM019** | Δ*smc::scar, attBφBT1*::pMS83-*smc-FLAG*. (modified TM010) ($Hyg^R$) | This study |

**Table S2 *E. coli* strains used in the study**

| Strain | Relevant genotype and characteristics | Source |
|---|---|---|
| DH5α | *F-, Φ80dlacZΔM15, recA1, endA1, gyrA96, thi-E1, hsdR17, (rk-,mk+), supE44, relA1, deoR, Δ(lacZYA-argF)U169* | Laboratory stock |
| BW25113/pIJ790 | (*araD-aarB*)567, Δ*lacZ*4787(::*rrnB*-4), *lacIp*-40000(*lacIQ*), λ-, *rpoS*369(*Am*), *rph-1*, Δ(*rhaD-rhaB*)568, *hsd*R514;, pIJ790 [*ori*R101], [*repA*1001(ts)], *araBp-gam-be-exo* | Laboratory stock [3] |
| ET12567/pUZ8002 | *dam, dcm, hsdS,*CmR, TetR, pUZ8002*: tra*, KanR, *RP4 23;* | Laboratory stock[4] |

**Table S3 Plasmids and cosmids used in the study**

| Plasmid/Cosmid | Relevant genotype and characteristics | Source |
|---|---|---|
| pGEM-T-Easy | $Amp^R$ | Promega |
| pIJ773 | pBluescript KS(+) deriviatve (*ori*pUC), *oriT* (RK2), FRT, ($Amp^R$, $Apr^R$) | [3] |
| pIJ10700 | pBluescript KS(+), *oriT*(RK2), miejsca FRT ($Hyg^R$) | [3] |
| pCP20 | pSC101 derivative, Rep101(Ts) *flp* ($Cm^R$ $Amp^R$) | [3] |
| pKF351 *ftsZ-ypet apra* | pIJ6902 derivative, $attB_{\Phi C31}$, *ftsZ-ypet* under the control of the native $p_{ftsZ}$ promoter ($Apr^R$) | [2] |
| pKF351 *ftsZ-ypet hyg* | pIJ6902 derivative, $attB_{\Phi C31}$, *ftsZ-ypet* under the control of the native $p_{ftsZ}$ promoter ($Hyg^R$) | [2] |
| pKF351 *ftsZ-ypet spec* | pIJ6902 derivative, $attB_{\Phi C31}$, *ftsZ-ypet* under the control of the native $p_{ftsZ}$ promoter $Spec^R$ | [2] |
| pSS170 | pMS82[5] derivative, intergrative vector ($attB_{\Phi BT1}$)($Hyg^R$ ) | Kind gift from dr S. Schlimpert, John Innes Centre, Norwich, UK)[6] |
| pSS172 | pMS82[5] derivative, intergrative vector ($attB_{\Phi BT1}$ ) ($Hyg^R$ ), *hupA-mcherry* under the control of the native $p_{hupA}$ promoter ($Hyg^R$) | Kind gift from dr S. Schlimpert, John Innes Centre, Norwich, UK)[6] |
| pSS170-*FLAG* | pSS170 derivative ($attB_{\Phi BT1}$) *FLAG*, ($Hyg^R$) | This study |
| pSS170-*hupS-FLAG* | pSS170 derivative ($attB_{\Phi BT1}$) *hupS-FLAG* under the control of the native $p_{hupS}$ promoter ($Hyg^R$) | This study |

| | | |
|---|---|---|
| Sv-3-B07 | Cosmid (SuperCos-1) that contains *S. venezuelae* chromosomal fragment encompasing *smc* gene (*vnz_26075*) (Amp$^R$, Kan$^R$) | Kind gift from prof. Mark Buttner, John Innes Centre, Norwich, UK) |
| Sv-5-D08 | Cosmid (SuperCos-1) that contains *S. venezuelae* chromosomal fragment encompasing *hupS* gene (*vnz_25950*) (Amp$^R$, Kan$^R$) | Kind gift from prof. Mark Buttner, John Innes Centre, Norwich, UK) |
| Sv-4-A09 ΔparB::apra | Cosmid (SuperCos-1) Δ*parB:: apra-oriT* (Amp$^R$, Kan$^R$) | [2] |
| Sv-3-B07 Δ*smc::apra* | Cosmid Sv-3-B07 Δ*smc:: apra-oriT* (Amp$^R$, Kan$^R$, Apr$^R$) | This study |
| Sv-3-B07 Δ*smc::scar* | Cosmid Sv-3-B07 Δ*smc*::scar (Amp$^R$, Kan$^R$) | This study |
| Sv-3-B07 Δ*smc::scar* (Apr$^R$) | Cosmid Sv-3-B07 Δ*smc*::scar, *bla*::*apra-oriT* (Apr$^R$, Kan$^R$) | This study |
| Sv-3-B07 *apra-oriT-FLAG-smc* | Cosmid Sv-3-B07 *apra-oriT-FLAG::smc* (Amp$^R$, Kan$^R$, Apr$^R$) | This study |
| Sv-3-B07 *FLAG-smc* | Cosmid Sv-3-B07 *smc:: FLAG-smc* (Amp$^R$, Kan$^R$) | This study |
| Sv-3-B07 *FLAG-smc* (Apr$^R$) | Cosmid Sv-3-B07 *smc:: FLAG-smc, bla*:: *apra-oriT* (Apr$^R$, Kan$^R$) | This study |
| Sv-5-D08Δhup*S::apra* | Cosmid Sv-5-D08 Δ*hupS*::*apra-oriT* (Amp$^R$, Kan$^R$, Apr$^R$) | This study |

**Table S4 Oligonucleotides used in the study**

| Primer | sequence |
|---|---|
| $p_{blaP1}$ | AATCTAAAGTATATATGAGTAAACTTGGTCTGACAGTTATGTAGGCTGGAGCTGCTTC |
| $p_{blaP2}$ | CCCTGATAAATGCTTCAATAATATTGAAAAAGGAAGAGTATTCCGGGGATCCGTCGACC |
| $p_{smc\_Fw}$ | GGCAAGTCCAACGTCGTGGACGCCCTCTCCTGGGTCATGCATATGATTCCGGGGATCCGTCGACC |
| $p_{smc\_Rv}$ | GGGTTCAACACTTGAAGCAATGGGGCATGCCCGGCCTCACATATGTGTAGGCTGGAGCTGCTTC |
| $p_{hupS\_Fw}$ | ACGGTACCCATATGATTCCGGGGATCCGTCG |
| $p_{hupS\_Rw}$ | ACGAATTCCTTGTCATCGTCATCCTTGTAATCGATGTCATGATCTTTATAATCACCGTCATGGTCTTTGTAGTCCATATGTGTAGGCTGGAGCTG |
| $p_{apra\_FLAG\_Rv}$ | ACGAATTCCTTGTCATCGTCATCCTTGTAATCGATGTCATGATCTTTATAATCACCGTCATGGTCTTTGTAGTCCATATGTGTAGGCTGGAGCTG |
| $p_{apra\_FLAG\_Fw}$ | ACGGTACCCATATGATTCCGGGGATCCGTCG |
| $p_{sv\_smc\_spr\_Fw}$ | CGCCGTTCCCCTCGTGTCC |
| $p_{sv\_smc\_spr\_rv}$ | CCTTCAAGTTTCGAAGTCAATACC |
| $p_{hupSspr\_Fw}$ | CAGGAACTCCGCAGCGGATCT |
| $p_{hupSspr\_Rv}$ | CGGATCACCCTGGTGACGCTC |
| $p_{ftszypet\_fw}$ | GCGGCCTTTGACTCCCTGC |
| $p_{ftszypet\_Rv}$ | CCATCTCCGGCGGCAGCG |

| | |
|---|---|
| $P_{pSS170\_xhoI\_FLAG\_Fw}$ | AGCTCTCGAGGACTACAAAGACCATGACG |
| $P_{pSS170-eco32I\_FLAG\_Rv}$ | CCAAGCTGATATCGAATTCGTAATCATGTCATAG |
| $P_{pSS170\_hupS\_FLAG\_Fw}$ | AGCTGGTACCACCGGTTGATGAAGGACCTCGACGAGGGC |
| $P_{hupS-tocherry-Rv}$ | CCAAGCTGATATCGAATTCGTAATCATGTCATAG |
| $P_{pSS\_spr\_Fw}$ | TTACCTCGCCTCTGACCCCTG |
| $P_{pSS\_spr\_Rv}$ | GGTTCATGTGCAGCTCCATCAGC |
| $p_{smc\_promoter\_Fw}$ | AGCTGGTACCCACGCTCCGACACCGGCC |
| $p_{smc\_NFLAG\_Rv}$ | CCGCGGAGGGTCAGGGCCTTGAGGTGCACCTTGTCATCGTCATCCTTGTAATCGATGT C |
| $p_{gyr2\_Fd}$ | GCTCCGCTATCACAAGATCA |
| $p_{gyr2\_Rv}$ | ACAGGAAGGTCAGCAGCAG |
| $p_{garg3\_Fd}$ | ACAGGAAGGTCAGCAGCAG |
| $p_{garg3\_Rv}$ | CCACTCCGACATCTCCTTG |
| $p_{smc\_poczatek\_Fw}$ | AGCTCCATGGCATATGGTGCACCTCAAGGCCCTGACCCTCCGC |
| $p_{smc\_exp\_Rv}$ | AGCTGGATCCCCGCGCAGCCGCTGGCTGATCAC |

**Construction of *smc* mutant strain (TM010)**

To construct unmarked *smc* deletion *S. venezueale* strain, first apramycine resistance cassette was inserted into the chromosome to replace *smc* gene using λ Red recombination based PCR-targeting protocol[3]. First the cassette encompassing *apra* gene and *oriT* was amplified on the template of pIJ773 using primers $p_{smc\_Fw}$ and $p_{smc\_Rv}$ and used to transform BW25113 carrying cosmid Sv-3-B07 and λ RED plasmid, pIJ790. The resulting cosmid Sv-3-B07Δ*smc*::*apra* was used for conjugation into wild type *S. venezuelae*. The $Apr^R$ exconjugants were screened for the loss of $Kan^R$, indicating a double-crossover allelic exchange of the *smc* locus. The obtained colonies were verified by sequencing of the PCR products obtained using the chromosomal DNA as the template and the primers $p_{sv\_smc\_spr\_Fw}$ and $p_{sv\_smc\_spr\_Rv}$. The obtained strain TM001 was next used for further modifications.

To obtain unmarked *smc* deletion strain, FLP recombinase, that recognises FRT sites flanking *apra* cassette, was used. To this end, DH5α containing pCP20 plasmid, that encodes yeast FLP recombinase, were transformed with the cosmid Sv-3-B07 *Δsmc::apra*. $Kan^R$ and $Apr^R$ resistant transformants were incubated at 42°C and next clones sensitive to apramycin were selected. After verification of the *scar* sequence generated in *smc* locus, the obtained Sv-3-B07 *Δsmc::scar* cosmid was modified, using PCR-targeting protocol, by insertion of *apra-oriT* cassette in the *bla* locus in the SuperCos. The cassette was amplified at the template of pIJ773 plasmid using primers $p_{blaP1}$ and $p_{blaP2}$. The resulting cosmid Sv-3-B07 *Δsmc::scar* was used for conjugation into *S. venezuelae smc::apra* (TM001) strain. The $Apr^R$, $Kan^R$ exconjugats were next screened for the loss of $Kan^R$ $Apr^R$, indicating a double-crossover allelic exchange of the *smc* locus. The obtained strain TM010 was verified by sequencing of the PCR products obtained using the chromosomal DNA as the template and the primers $p_{sv\_smc\_spr\_Fw}$ and $p_{sv\_smc\_spr\_Rv}$.

**Construction of *FLAG-smc* (TM017) and *parB FLAG-smc* (KP4F4) mutant strains**

Strains producing FLAG-SMC were constructed by homologous recombination and modification of *smc* gene in its native chromosomal locus. First the apramycin resistance cassette flanked with the oligonucleotide encoding FLAG was amplified on the template of pIJ773, using $p_{apra\_FLAG\_Fw}$ and $p_{apra\_FLAG\_Rv}$ primers that contained NdeI restriction sites and used to transform BW25113 carrying cosmid Sv-3-B07 and λ RED plasmid, pIJ790. Next, the obtained cosmid Sv-3-B07 *apra-oriT- FLAG-smc* was digested with NdeI and religated to remove *apra-oriT* from the upstream region of *smc* gene. The obtained cosmid was verified using PCR with primers $p_{smc\_promoter\_Fw}$ and $p_{smc\_NFLAG\_Rv}$ and then modified, using PCR-targeting protocol, by insertion of *apra-oriT* cassette in the *bla* locus in the SuperCos. The cassette was amplified at the template of pIJ773 plasmid using primers $p_{blaP1}$ and $p_{blaP2.}$ The resulting disrupted cosmid Sv-3-B07 *FLAG-smc* was used for conjugation into *S. venezuelae smc::apra* (TM001) strain. The $Apr^R$, $Kan^R$ exconjugates were next screened for the loss of $Kan^R$ $Apr^R$, indicating a double-crossover allelic exchange of the *smc* locus. The obtained strain TM017 was verified by sequencing of the PCR products obtained with the primers $p_{smc\_promoter\_Fw}$ and $p_{smc\_NFLAG\_Rv}$ on the template of the chromosomal DNA as well as Western Blotting with the anti-FLAG M2 antibody (Merck).

To obtain *parB FLAG-smc* strain (KP4F4) cosmid Sv-4-A98Δ*parB::apra* was used for conjugation into *S. venezuelae FLAG-smc* (TM017) strain. The $Apr^R$ exconjugants were screened for the loss of $Kan^R$, indicating a double-crossover allelic exchange of the *parAB* locus.

**Construction of *hupS* (AKO200), *hupS smc* (TM003) and *hupS FLAG-smc* (TM018) mutant strains**

*S. venezuelae hupS* deletion strain was constructed using λ Red recombination PCR-targeting method[3] by introduction of *apra* resistance cassette in the *hupS* locus. First, the cassette encompassing *apra* gene and *oriT* was amplified on the template of pIJ773 using primers $p_{hupS\_Fw}$ and $p_{hupS\_Fw}$ and used to transform BW25113 carrying cosmid Sv-5-D08 and λ RED plasmid, pIJ790. The resulting cosmid Sv-5-D08Δ*hupS*::*apra* was used for conjugation into the wild type, *smc::scar* (TM010) and to *FLAG-smc* (TM017) strains. The $Apr^R$ exconjugants were screened for the loss of $Kan^R$, indicating a double-crossover allelic exchange in the *hupS* locus. The obtained *hupS* (AKO200), *hupS smc* (TM003) and *hupS FLAG-smc* (TM018) strains were verified by sequencing of the PCR products obtained using the chromosomal DNA as the template and the primers $p_{hupSspr\_Fw}$ and $p_{hupSspr\_Rv}$.

**Construction of *hupS*-FLAG (TM015) and *hupS*-FLAG *smc* (TM016) mutant strains**

To construct *hupS*-FLAG strains plasmid pSS170-*hupS-FLAG,* that contained *hupS* gene under the control the native *hupS* promoter, was used. The oligonucleotide encoding FLAG was amplified using primers $P_{\text{pSS170\_xhoI\_FLAG\_Fw}}$ and $P_{\text{pSS170-eco32I\_FLAG\_Rv}}$ that included restriction sites XhoI and Eco32I and cloned into pSS170 plasmid (Hyg$^R$, kind gift from dr. Susan Schlimpert, John Innes Centre, Norwich, UK) delivering pSS170-*FLAG*. *hupS* gene with its native promoter was amplified on the template of cosmid Sv-5-D08 using primers $P_{\text{pSS170\_hupS\_FLAG\_Fw}}$ and $P_{\text{hupS-tocherry-Rv.}}$ The fragment containing *hupS* gene was cloned to pSS170-*FLAG* using restriction sites Acc65I and XhoI. The obtained plasmid pSS170-*hupS-FLAG* was used for conjugation to *hupS* (AKO200) and *hupS smc* (TM003) strains delivering *hupS-FLAG* (TM015) and *hupS-FLAG smc* (TM016). Strains were verified by sequencing of the PCR products obtained using primers $p_{\text{pSS\_spr\_Fw}}$ and $p_{\text{pSS\_spr\_Rv}}$ on the template of chromosomal DNA, and Western Blotting with anti-FLAG M2 antibody (Merck).

**Construction of *smc* and *hupS* mutant derivatives producing FtsZ-Ypet and HupA-mCherry**

To construct the *S. venezuelae* strains expressing *ftsZ-ypet* the obtained strains TM010, AKO200 and TM003 were used for conjugation with the *E. coli* carrying pKF351 *ftsZ-ypet apra* (to TM010) or pKF351 *ftsZ-ypet hyg* (to AKO200 and TM003) plasmid. *smc ftsZ-ypet* (TM004 Apr$^R$), *hupS ftsZ-ypet* (TM005, Hyg$^R$) and *hupS smc ftsZ-ypet* (TM006, Hyg$^R$) were verified by PCR with $p_{\text{ftszypet\_fw}}$ and $p_{\text{ftszypet\_Rv}}$ and fluorescence microscopy.

To construct the *S. venezuelae* strains expressing *ftsZ-ypet* and *hupA-mcherry,* we used plasmid pSS172*hupA-mcherry* (hyg$^R$, kind gift from dr. Susan Schlimpert, John Innes Centre, Norwich, UK) that contained *hupA-mCherry* under the control of the native *hupA* promoter. Plasmid *pSS172hupA-mCherry* was introduced to *ftsZ-ypet* (in the wild type background, MD100, Apr$^R$) and *smc ftsZ-ypet* (TM004, Apr$^R$) strain generating *ftsZ-ypet hupA-mcherry* (TM011, Apr$^{R,}$ Hyg$^R$) and *smc ftsZ-ypet hupA-mCherry* (TM012, Apr$^{R,}$ Hyg$^R$). Strains *hupS ftsZ-ypet*, *hupA-mCherry* (TM013, Apr$^{R,}$ Hyg$^R$ Spec$^R$) and *hupS smc ftsZ-ypet hupA-mCherry* (TM014, Apr$^{R,}$ Hyg$^R$ Spec$^R$) were constructed by subsequent delivery of pKF351 *ftsZ-ypet spec* and pSS172*hupA-mCherry* to *hupS* (AKO200) and *hupS smc* (TM003). The obtained strains were verified by PCR with $P_{\text{pSS\_spr\_Fw}}$ and $P_{\text{pSS\_spr\_Rv}}$ primers using chromosomal DNA as template and by using fluorescence microscopy.

**Construction of *smc* complemented with *smc-FLAG***

Genetic complementation of Δ*smc::scar* (TM010) was performed by delivering *smc-FLAG in trans*. To this end the *smc* gene was amplified using cosmid DNA Sv-3B07 as template, using primers $p_{\text{smc\_poczatek\_Fw}}$ and $p_{\text{smc\_exp\_Rv}}$, next PCR product was cloned into pGEM-T-Easy vector, to which also double stranded oligonucleotide encoding FLAG was inserted using BamHI and Mph1103I restriction enzymes. The promoter region of *smc* was amplified using Sv-3-B07 comid DNA as template and primers $p_{\text{smc\_promotoer\_Fw}}$ and $p_{\text{smc\_promoter\_Rv}}$ and cloned directly into pMS83 using KpnI and NdeI. Next, fragment encoding SMC-FLAG was cloned from pGEM-TEasy-*smc-FLAG* to pMS83-$p_{smc}$ yielding pMS83-

*smc-FLAG,* which was used for conjugation to *Δsmc::scar* (TM010) yielding TM019 (*Δsmc::scar*; pMS83-*smc-FLAG*). The construct was verified by sequencing of PCR products obtained using the chromosomal TM019 as template and primers $p_{smc\_spr\_Fw}$ and $p_{smc\_spr\_Rv;}$ as well as $p_{smc\_inter\_Fw}$ and $p_{smc\_NFLAG\_Rv,.}$ and then by Western blotting.


1. Bibb, M. J., Domonkos, A., Chandra, G. & Buttner, M. J. Expression of the chaplin and rodlin hydrophobic sheath proteins in Streptomyces venezuelae is controlled by σ(BldN) and a cognate anti-sigma factor, RsbN. *Mol. Microbiol.* **84**, 1033–1049 (2012).
2. Donczew, M. *et al.* ParA and ParB coordinate chromosome segregation with cell elongation and division during Streptomyces sporulation. *Open Biol.* **6**, 150263 (2016).
3. Gust, B. *et al.* Red-Mediated Genetic Manipulation of Antibiotic-Producing Streptomyces. *Adances Appl. Microbiol.* **54**, 107–128 (2004).
4. Kieser, T., Bibb, M. J., Buttner, M. J., Chater, K. F. & Hopwood, D. A. Practical Streptomyces Genetics. *John Innes Cent. Ltd.* 529 (2000) doi:10.4016/28481.01.
5. Gregory, M. A., Till, R. & Smith, M. C. M. Integration Site for Streptomyces Phage [illegible] BT1 and Development of Site-Specific Integrating Vectors. *J. Bacteriol.* **185**, 5320–5323 (2003).
6. Ramos-léon, F., Bush, M. J., Sallmen, J. W. & Chandra, G. A conserved cell division protein directly regulates FtsZ dynamics in filamentous and unicellular actinobacteria. *BioRxive* (2020).
